# pChem: a modification-centric assessment tool for the performance of chemoproteomic probes

**DOI:** 10.1101/2021.09.22.461295

**Authors:** Ji-Xiang He, Zheng-Cong Fei, Ling Fu, Cai-Ping Tian, Fu-Chu He, Hao Chi, Jing Yang

**Affiliations:** College of Chemistry & Environmental Science, Hebei University, Baoding, 071002, China; State Key Laboratory of Proteomics; Beijing Proteome Research Center; National Center for Protein Sciences • Beijing; Beijing Institute of Lifeomics, Beijing, China; Key Laboratory of Intelligent Information Processing of Chinese Academy of Sciences (CAS); Institute of Computing Technology, CAS; University of Chinese Academy of Sciences, Beijing, China

## Abstract

Chemoproteomics has emerged as a key technology to expand the functional space in complex proteomes for probing fundamental biology and for discovering new small molecule-based therapies. Here we report a modification-centric computational tool termed <u>pChem</u> to provide a streamlined pipeline for unbiased performance assessment of chemoproteomic probes. The pipeline starts with an experimental setting for isotopically coding probe-derived modifications (PDMs) that can be automatically recognized by pChem, with masses accurately calculated and sites precisely localized. Further, pChem exports on-demand reports by scoring the profiling efficiency, modification-homogeneity and proteome-wide residue selectivity of a tested probe. The performance and robustness of pChem were benchmarked by applying it to eighteen bioorthogonal probes. Of note, the analyses reveal that the formation of unexpected PDMs can be driven by endogenous reactive metabolites (e.g., bioactive aldehydes and glutathione). Together, pChem is a powerful and user-friendly tool that aims to facilitate the development of probes for the ever-growing field of chemoproteomics.

---

Chemical probe coupled with mass spectrometry (MS)-based proteomics, herein termed chemoproteomics, offers versatile tools to globally profile protein features and to systematically interrogate the mode of action of small molecules in a native biological system^1^. For instance, bioorthogonal probes surrogating endogenous metabolites (e.g., sugars and lipids) enable the proteome-wide mapping of post-translational modifications (PTMs) on specific amino acid residues^2^. In addition, various activity-based protein profiling (ABPP) probes have been developed by targeting amino acid residues including cysteine^3, 4^, lysine^5^, tyrosine^6^, methionine^7, 8^, histidine^9^, aspartate and glutamate^10, 11^, as well as their PTM forms^12–14^, which greatly expand the chemical space in complex proteomes for probing fundamental biology and for discovering new small molecule-based therapies.

Nonetheless, the development of an efficient and selective probe for chemoproteomics can still be challenging. It is particularly difficult to unbiasedly assess its chemoselectivity at a proteome-wide scale, since a chemical probe displaying selectivity well-characterized *in vitro* would possibly generate unexpected modifications owing to potential cross-reactivity in complex biological systems. In addition, unforeseeable probe-derived modifications (PDMs) may be yielded during sample preparation or in-source MS fragmentation, thereby causing inhomogeneous modifications on the same sites and complicating data analysis.

Notably, the last decade has witnessed tremendous progress in the development of blind search informatic tools^15–23^. Such tools, in combination with isotope-coding approaches for probes (**Supplementary Fig. 1**), can provide an unbiased survey of PDMs that can be distinguished from non-probe-derived modifications (e.g., unmodified or endogenously modified ones), considering that only those peptides bearing PDMs would yield an isotopic MS signature (**Fig. 1a**). For example, we have previously used this pipeline (i.e., TagRecon for blind search^23^) to substantiate the performance of several newly developed chemoselective probes for proteomic mapping of cysteine redox forms^12, 13^. The pipeline also allowed us to uncover unexpected PTMs being captured by chemoproteomic probes *in situ*^14, 24^. Most recently, Hacker and coworkers have applied a similar pipeline (i.e., MSFragger for blind search^18^) to systematically investigate the proteome-wide selectivity of diverse electrophilic probes^25^. These studies underscore the power of blind search tools, which provide an ideal means to unbiasedly assess the proteome-wide residue selectivity of a probe and to uncover new chemotypes in the proteome.

**Fig. 1.**
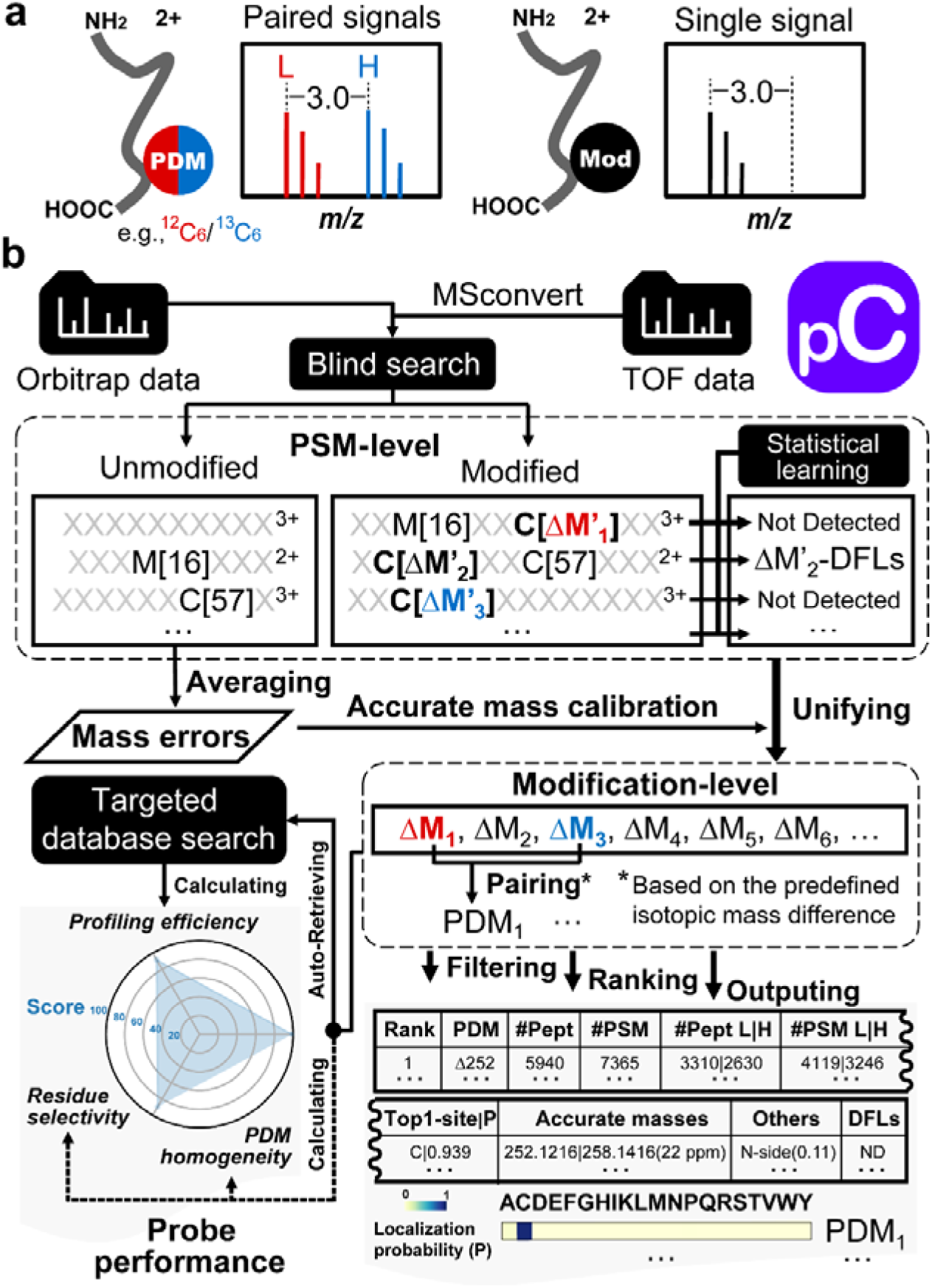
Principle and workflow of pChem. **a**, Isotope-coding of probe-derived modifications (PDMs) yields a paired MS signature (Left) that can be distinguished from non-PDMs (Right, i.e., endogenous and/or common modifications). **b**, Schematic of pChem workflow. MS data is first imported into pChem to perform *bona fide* blind search against the protein sequence database, which generates abundant modification candidates at the PSM level. Meanwhile, PDM-specific diagnostic fragment losses (DFLs) are recognized with a statistic learning approach. Note that the candidates allow modification (ΔM’) incorporated from the common modifications (e.g., methionine oxidation Δ16 and cysteine carbamidomethylation Δ57) that can be manually or automatically pre-defined. After accurate mass calibration, those modification candidates with the same calculated ΔM’ from multiple PSMs are unified together and paired according to the predefined isotopic mass difference. Then, several semi-empirical criteria (See **Methods**) are applied to automatically define the high-confidence PDMs and ranked based on abundance. All on-demand information for probe developers is summarized in an output table and heatmaps. Optionally, a radar plot can be generated for illustrating overall performance outcomes by scoring the profiling efficiency (via an additional round of targeted database search automatically invoked by pChem), modification-homogeneity and residue selectivity.

Despite these advances, a set of challenges have emerged. First, most blind search tools cannot automatically unify the localization probability (or residue selectivity) and accurate masses of PDMs, as most of them only offer identification and site localization at a PSM (peptide-spectrum match) level. Second, no available tools can automatically distinguish isotopically coded PDMs from non-probed ones, while manual evaluation of dozens to hundreds of mass shifts can be a daunting task. Last but not the least, for probe developers, even those with substantial bioinformatics expertise, managing the existing tools can be tedious as all require laborious installation and setup, and output many redundant information rather than on-demand reports.

The challenges discussed above have therefore inspired us to develop an automated, user-friendly, fit-for-purpose computational tool for the ever-growing field of chemoproteomics. Here we present pChem, a modification-centric blind search and summarization tool to provide a pipeline for rapid and unbiased assessing of the performance of ABPP and metabolic labeling probes. This pipeline starts experimentally by isotopic coding of PDMs, which can be automatically recognized, paired, and accurately reported by pChem, further allowing users to score the profiling efficiency, modification-homogeneity and proteome-wide residue selectivity of chemoproteomic probe.

## Results

### The design of pChem

Building upon the <u>pFind</u> platform^15^, pChem first aims to generate all possible modifications using *bona fide* blind search based on a tag-index approach (**Fig. 1b**). Specifically, for each spectrum, a number of 5-mer sequence tags are extracted with a depth-first *de novo* search. The resulting tags are then scored and up to 100 ranked tags are retained before searching against the tag-indexed protein database in order to extend to a full-length peptide sequence. During the extension procedure, all mass shifts within the given range (±1,000 Da by default) are considered as modification candidates. Note that the default-defined common modifications (e.g., oxidation of methionines and carbamidomethylation of cysteines) are involved in the sequence tag extraction procedure to improve the sensitivity of this algorithm. Alternatively, a few highly abundant modifications can be automatically detected by the initial pFind-based open-search against Unimod^26^, a protein modification database for mass spectrometry applications, and then be specified in the following blind search.

Similar to other blind search tools, pChem will initially generate numerous mass shifts at the PSM-level, and then the mass shifts can be calibrated with the system error computed from PSMs identified without any unknown modifications. Specifically, the system error of the precursor ion mass is obtained by averaging each mass difference between the precursor ion mass and the theoretical mass of the peptide without any modifications or with known modifications. The calibrated mass shifts on PSMs are further grouped together and the mean value is calculated for each group in order to generate the accurate mass for modification candidates and score their residue-specific localization probability (**Fig. 1b**). To eliminate non-PDMs or other unrealistic modifications, all modification-centric mass shifts are automatically paired according to the isotopic mass difference (e.g., 6.020132 Da caused by six heavy carbons) within empirically defined tolerance (i.e., ±0.001 Da for a fine-tuned Orbitrap instrument). Furthermore, any trace PDM (i.e., byproducts from an unwanted side reaction or rearrangement) with the percent of total PSMs less than 5% by default can be neglected and therefore not reported (See **Methods**). Meanwhile, using an unsupervised outlier-detection approach based on statistical learning, pChem automatically recognizes and reports diagnostic fragment losses (DFLs, e.g., neutral and/or charged losses), if any, which facilitates the characterization of each PDM.

Finally, the pChem results will be exported into the heatmaps showing the site localization probability of all high-confidence PDMs and into several editable files for users to explore more details. Optionally, by automatic retrieving the accurate PDMs for the final round of targeted database search (i.e., a restricted search procedure eliminating the lengthy and laborious process of re-setting and re-searching modifications deduced from numerous mass shifts), pChem can also generate a radar plot showing overall performance outcomes by scoring the profiling efficiency, modification-homogeneity, and residue selectivity (**Fig. 1b**). Note that pChem is an installation-free executable (.exe) program that can be easily used on either laptops or desktops with the step-by-step guidance for operation and troubleshooting (**Supplementary Protocol**).

### Performance and robustness of pChem

To benchmark the performance of pChem, we initially generated a QTRP (quantitative thiol reactivity profiling) dataset by using a ‘clickable’ iodoacetamide probe termed IPM (**Fig. 2a**) with a well-established chemoselectivity towards cysteinyl thiols^4^. Specifically, the HEK293T cell lysates were labeled with IPM and then processed into a mixture of isotopically tagged peptides with a predefined light to heavy ratio of 1.0 via CuAAC (Cu^I^-catalyzed azide-alkyne cycloaddition, one type of click chemistry reactions)-mediated biotin-conjugation. The isotopically modified peptides were captured with streptavidin and selectively released for MS analysis on a Q-Exactive Plus instrument. For each of three biological replicates, pChem reports the targeted PDM of Δ252.12 Da (# PSM counts: 7762 ± 448) on cysteine with a high localization probability (94.7 ± 1.2%, **Fig. 2b**)^27^ and a minor one (Δ268.12 Da, # PSM counts: 713 ± 86) that is mainly attributed to the inevitable mis-matching due to an oxidized methionine adjacent to the cysteine bearing Δ252.12 (**Supplementary Fig. 2**, **Supplementary Table 1-2**, and **Supplementary Note 1**). Moreover, pChem scored high on the overall performance of the IPM-based chemoproteomic method (**Fig. 2c**).

**Fig. 2.**
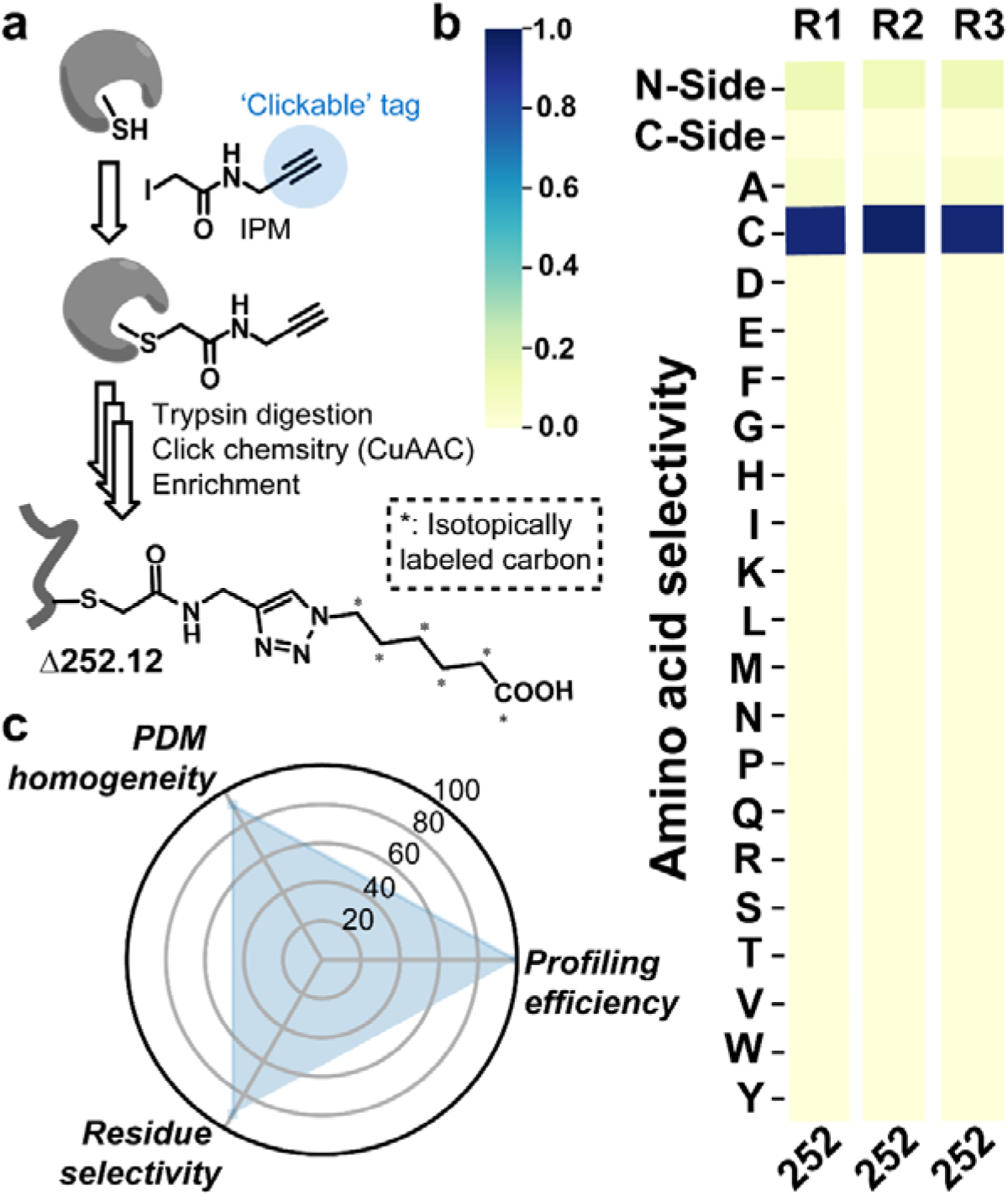
Performance of pChem. **a**, Chemical structures of IPM and its expected PDM. **b**, Heatmaps generated by the default pChem search (n=3) showing the amino acid localization distribution of the IPM-derived modification Δ252.12. **c**, A representative radar plot showing the overall performance of IPM for chemoproteomic profiling of the cysteinome.

To validate the robustness of pChem, we generated additional QTRP samples and analyzed them on eight different instruments from three vendors. Note that the *\*.RAW* data generated by Thermo Scientific*™* Orbitrap instruments can be directly imported into pChem, while the *\*.WIFF* (SCIEX) and *\*.TDF* (Bruker) data from Time-of-Flight (TOF) instruments need to be first transformed into *\*.mzML* (mass spectrometry markup language) using MSconvert^28^. As expected, the results as above could be reproduced (**Supplementary Fig. 3a** and **Supplementary Table 3**), demonstrating pChem as a cross-platform tool. Next, we sought to evaluate the effect of predefined light to heavy ratios on the performance of pChem. To this end, we analyzed a published QTRP dataset from light and heavy tagged samples mixed in different ratios (L/H = 1:10, 1:5, 1:2, 1:1, 2:1, 5:1 and 10:1) that has been created for quantification accuracy evaluation^4^. Also, pChem generated the same results, no matter how one pre-mixed the light and heavy tagged samples (**Supplementary Fig. 3b** and **Supplementary Table 3**). Likewise, we found that pChem analysis was insensitive to the types of samples/species and the corresponding protein databases (**Supplementary Fig. 3c** and **Supplementary Table 2**). Taken together, the results above demonstrate pChem as a robust assessment tool for the performance of chemoproteomic probes.

### Application of pChem for residue-reactive probes

To further evaluate the performance of pChem, we applied it to re-assess several residue-reactive probes in their usage for chemoproteomics.

First, we used pChem to analyze a set of newly collected datasets generated by using various covalent probes with diverse electrophilic warheads (**Fig. 3a**), including four cysteine-reactive probes (incl. ENE, NPM, PPMS and VSF) and two lysine-reactive probes (incl. NHS and STP). Notably, the analyses not only unbiasedly identified the targeted PDM corresponding to each probe (**Fig. 3b**, **Supplementary Tables 1 and 4**), but also explained the prevalence of selected warheads for cysteine or lysine bioconjugation (**Fig. 3c**). Meanwhile, pChem allowed us to uncover a previously unknown N-term cysteine modification (Δ292.12) derived from the NPM probe (**Fig. 3b** and **Supplementary Fig. 4**). Interestingly, this modification predominantly occurred at the carboxyl side of lysine or arginine on proteins, suggesting that it formed after tryptic digestion. As shown in **Fig. 3d**, a mechanism was proposed as follows: **a**) NPM rapidly forms Michael adducts with protein cysteine residues. **b**) when peptide bonds are hydrolyzed on the carboxyl side of K/R adjacent to the initially adducted cysteine residue, the newly formed primary amine attacks the succinimido ring to undergo the intramolecular aminolysis^29^. We also generated a dataset using a ‘cocktail’ of three thiol-reactive probes, including IPM, ENE, and NPM (**Supplementary Fig. 5a**). Notably, all and only the targeted PDMs could be automatically found by pChem (**Supplementary Fig. 5b** and **Supplementary Table 4**), which further validates the sensitivity and accuracy of this tool.

**Fig. 3.**
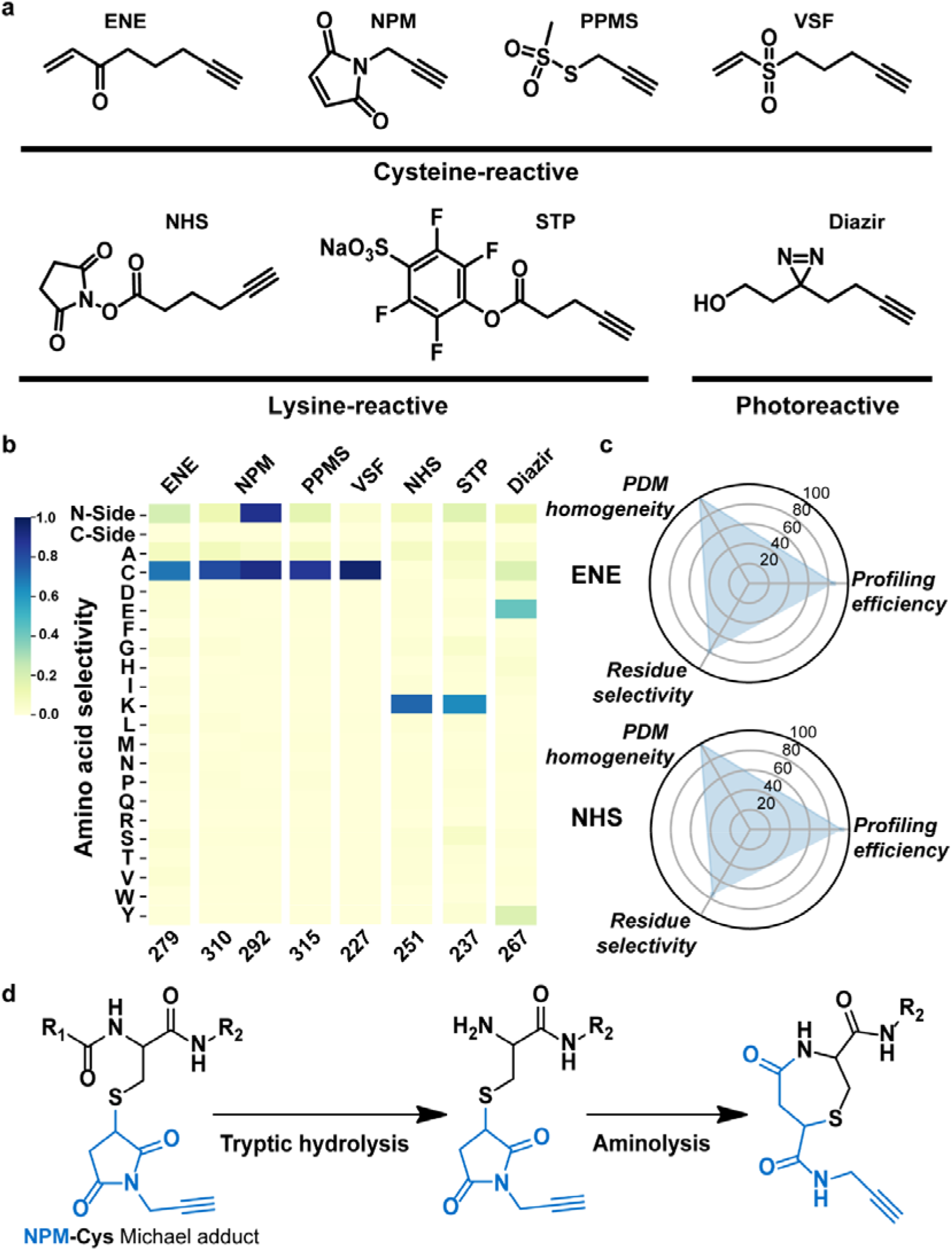
Application of pChem for residue-reactive probes. **a**, Chemical structures of the residue-reactive probes for benchmarking pChem. **b**, Representative heatmaps showing the amino acid localization distribution of the pChem-defined PDMs for each indicated probe. **c**, Representative radar plots showing the overall performance of ENE (Warhead: unsaturated ketone) and NHS (Warhead: activated ester) for cysteine and lysine profiling, respectively. **d**, Plausible mechanism for the formation of an unexpected N-term cysteine modification (Δ292.12) derived from NPM.

Next, we asked whether pChem could unbiasedly address the long-standing issue regarding the proteome-wide reactivity of the diazirine group that has been widely used in developing photoaffinity labeling probes^30^. Here we generated a dataset for pChem analysis from a proteome sample labeled with an alkyl diazirine probe (**Fig. 3a**). To our surprise, such a probe that has been considered lack of residue-selectivity preferentially react with glutamic acid (**Fig. 3b**, **Supplementary Fig. 6**, **Supplementary Tables 1 and 4**). Notably, during the preparation of this manuscript, a recent study also systematically uncovered the labeling preference of diazirine probes toward acidic amino acids^31^, in accordance with our observation to some extent.

Finally, we turned our attention to probes enabling enzymatic proximity labeling (**Supplementary Fig. 7a**), which has emerged as the method-of-choice to globally study protein-protein interactions in living biological systems^32^. For instance, phenol probes have been predominantly used in peroxidase-catalyzed proximity labeling^33^. By using restricted search, phenol probes have been shown to almost solely react with tyrosine (i.e., >98% of labeling sites) via live-cell APEX peroxidase-based proximity labeling^34, 35^. To further investigate the proteome-wide residue selectivity by peroxidase-mediated proximity labeling in a more unbiased fashion, we generated a dataset using two types of peroxidases (i.e., *in cellulo* APEX and *in vitro* HRP) with alkyne phenol (AP) as the labeling probe (**Supplementary Fig. 7b**). The pChem analysis then revealed that APEX-catalyzed probe labeling indeed predominantly occurred on tyrosine in an expected form (Δ372.18), while an unusual PDM of Δ209.08 was also mapped onto many lysine sites (**Supplementary Fig. 7c-d** and **Supplementary Tables 1 and 4**). The latter may be formed by the reaction between lysine and an acyl radical intermediate probably generated from the APEX-catalyzed radical reaction (**Supplementary Fig. 7e**)^36, 37^. Instead, HRP-mediated labeling sites were mapped on both tyrosine and cysteine (**Supplementary Fig. 7c**). It is worth noting that cysteine was modified by the AP probe via an *o*-quinone intermediate (**Supplementary Fig.7d-e** and **Supplementary Tables 1 and 4**). In fact, it has been historically known that HRP-catalyzed oxidation of phenols can produce quinonoid reactive species to adduct free thiol group^38, 39^. Regardless, this finding suggests that the selectivity of phenol probes for proximity labeling may vary in different enzymatic conditions.

### Application of pChem for oxoform-specific probes

Cysteine oxoforms, such as sulfenic acid (−SOH) and sulfinic acid (−SO_2_H), represent important post-translational approaches to regulate protein functions in a redox-dependent manner^40^. In the last decade, the development of various cysteine oxoform-specific probes has greatly advanced the field of thiol-based redox biology^41^. In particular, some of such probes designed for MS-based redox proteomics allowed us to site-specifically map and quantify hundreds to thousands of cysteine oxoforms in complex proteomes (**Fig. 4a**), thereby greatly expanding the cysteine redoxome^12, 42–45^. Nevertheless, the chemoselectivity of oxoform-specific probes has long been controversial, even for those classic ones (i.e., dimedone for SOH)^46, 47^. To address this controversy in an unbiased fashion, we used pChem to re-analyze the redox proteomic datasets generated in our laboratory.

**Fig. 4.**
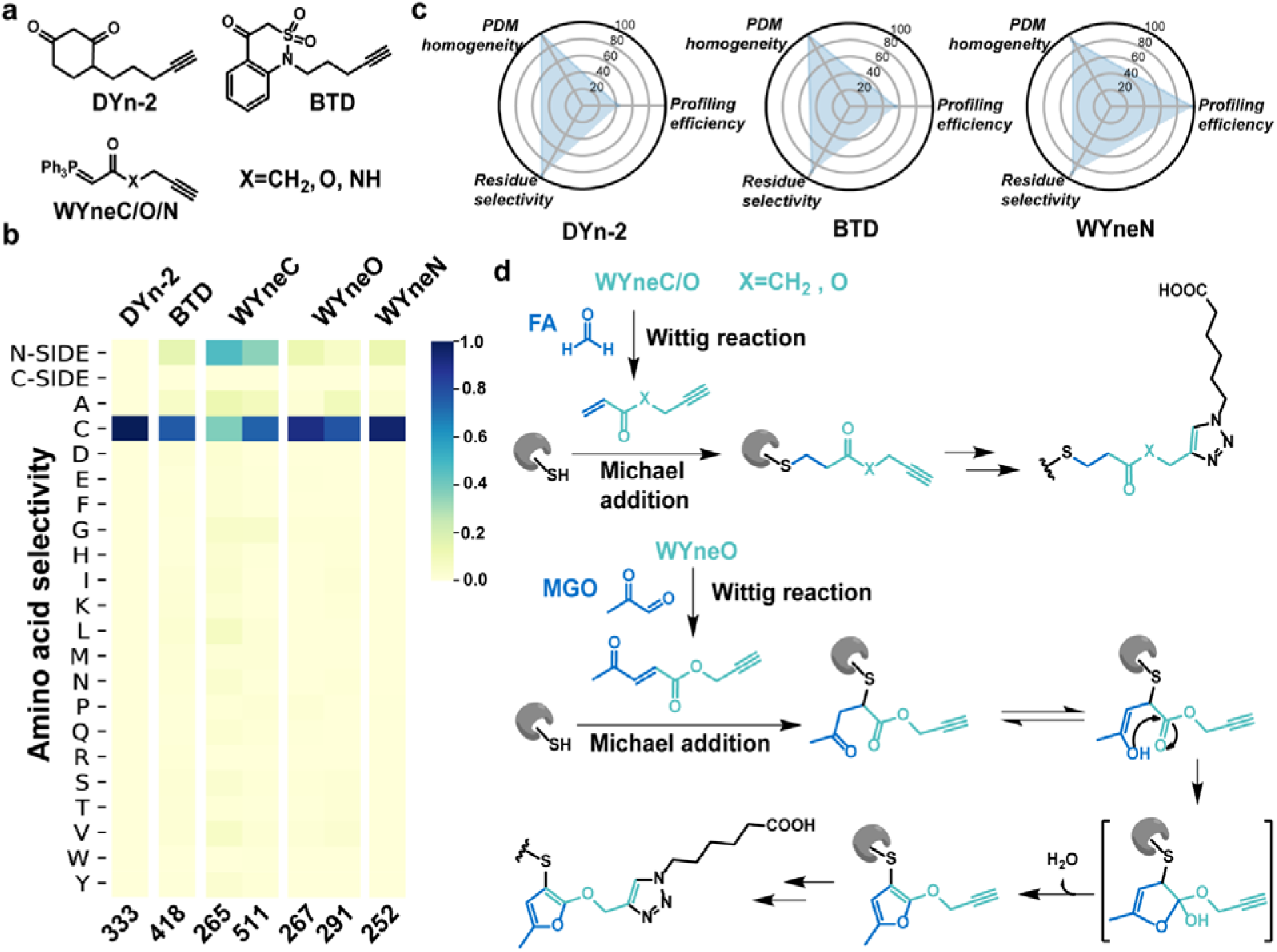
Application of pChem for SOH probes. **a**, Chemical structures of the SOH probes, including DYn-2, BTD, and WYneC/O/N. **b**, Representative heatmaps showing the amino acid localization distribution of the pChem-defined PDMs for each indicated probe. **c**, Representative radar plots showing the overall performance of the tested nucleophilic probes for profiling cysteine sulfenic acids. **d**, Plausible mechanisms for the formation of unwanted PDMs from WYneC/O.

As expected, the dimedone-based DYn-2 and the benzothiazine-based BTD, two most used probes for detecting and/or profiling protein SOHs^48, 49^, produced the corresponding targeted PDMs (i.e., Δ333.17 for DYn-2 and Δ418.13 for BTD) that were predominantly localized on cysteine (**Fig. 4b**, **Supplementary Tables 1 and 5**). As compared to DYn-2 and BTD, a phosphonium ylide-based SOH probe termed WYneN (**Fig. 4a**) that is reported most recently^50^ exhibited a superior overall performance (**Fig. 4b–c**). Notably, the pChem analysis revealed that the targeted PDM (Δ252.12), a triphenylphosphonium (TPP)-loss thioether product of the initial conjugate from WYneN showed a localization probability of >0.95 on cysteine. In the previous report, we also described two other two WYne probes (WYneC/O, **Fig. 4a**), albeit not ones for chemoproteomics due to the lack of profiling efficiency^50^. Note that WYneC/O were found to be able to produce TPP-loss PDMs as well, but they were originally assigned as thiol ester products with theoretical ΔMs of 265.1063 (C_12_H_15_N_3_O_4_) and 267.0855 (C_11_H_13_N_3_O_5_), respectively (**Supplementary Fig. 8** and **Supplementary Tables 1 and 5**). However, such a hypothesis was proven wrong according to the accurate masses calculated by pChem (Δ265.1441 for WYneC and Δ267.1213 for WYneO, **Fig. 4b** and **Supplementary Fig. 9**). Moreover, pChem revealed an additional PDM of Δ291.1207 for WYneO, which also seemed to be lack of TPP. As such, the formation of these TPP-loss products can be explained by the plausible mechanisms as follows (**Fig. 4d**). WYneC/O first react with endogenous formaldehyde and/or methylglyoxal (MGO, a by-product of glycolysis^51^) via the classic Wittig reaction to form α, β-unsaturated ketones followed by the Michael addition reaction of the latter with reduced cysteines or other biological nucleophiles. The new mechanism was further supported by the evidence from an *in vitro* labeling experiment^50^, in which we failed to detect any TPP-loss modifications from WYneC/O. Overall, this mechanism apparently precludes the use of WYneC/O for live-cell labeling protein sulfenic acids in any applications.

In addition, pChem confirmed that DiaAlk (**Supplementary Fig. 10a**) mostly targets protein sulfinic acids by generating the expected PDM of Δ387.17 (#PSMs: 172) on cysteine with the known neutral or charged losses (**Supplementary Fig. 10b**, **Supplementary Tables 1 and 5**)^12^. The analysis also provided two minor PDMs, including Δ471.23 (#PSMs: 46) and Δ369.17 (#PSMs: 12) both on tryptophan (**Supplementary Fig. 10b-d**), which are most likely formed from an unwanted side-reaction by DiaAlk. Specifically, the mechanism is initial attack at the electrophilic nitrogen of DiaAlk by C-2 of the tryptophan indole ring, followed by oxidation-induced ring open at C-3, resulting in an imine conjugate and its BOC (t-butyloxycarbonyl)-loss product (**Supplementary Fig. 10e**). One therefore needs to be cautious when using DiaAlk for any protein-centric analysis (*i.e.*, Western blotting), although this probe can still be applicable for site-centric chemoproteomic profiling of SO_2_H.

### Application of pChem for metabolite-derived probes

The advances in biorthogonal chemistry have inspired a series of ‘clickable’ metabolites as chemical reporters for various PTMs^2^. For instance, metabolic glycan labeling with the use of unnatural monosaccharides bearing an azide or alkyne reporter has become a widespread approach for studying protein glycosylation^52^. Recently, Qin et al.,^53^ reported a non-enzymatic reaction of an azido analog of N-azidoacetylmannosamine (Ac_4_ManNAz) with cysteine, which may interfere with its glycoproteomic profiling. pChem further confirmed this unwanted *S*-glycosylation by searching the chemoproteomic data from the Ac_4_ManNAz-labeling sample (**Supplementary Fig. 11**, **Supplementary Tables 1 and 6**). Encouraged by these efforts, we then extended pChem to re-analyze two public datasets that were generated from samples metabolically labeled by alkyne surrogates of endogenous lipid electrophiles, 4-hydroxy-2-nonenal (HNE) and 4-oxo-2-nonenal (ONE), which can readily modify cellular proteomes (**Fig. 5a**)^54^. Notably, our analyses not only reported almost all known PDMs by these probes^24, 55^, but also uncovered two major ones that have been overlooked in our original publications (**Fig. 5b–c**, **Supplementary Tables 1 and 6**).

**Fig. 5.**
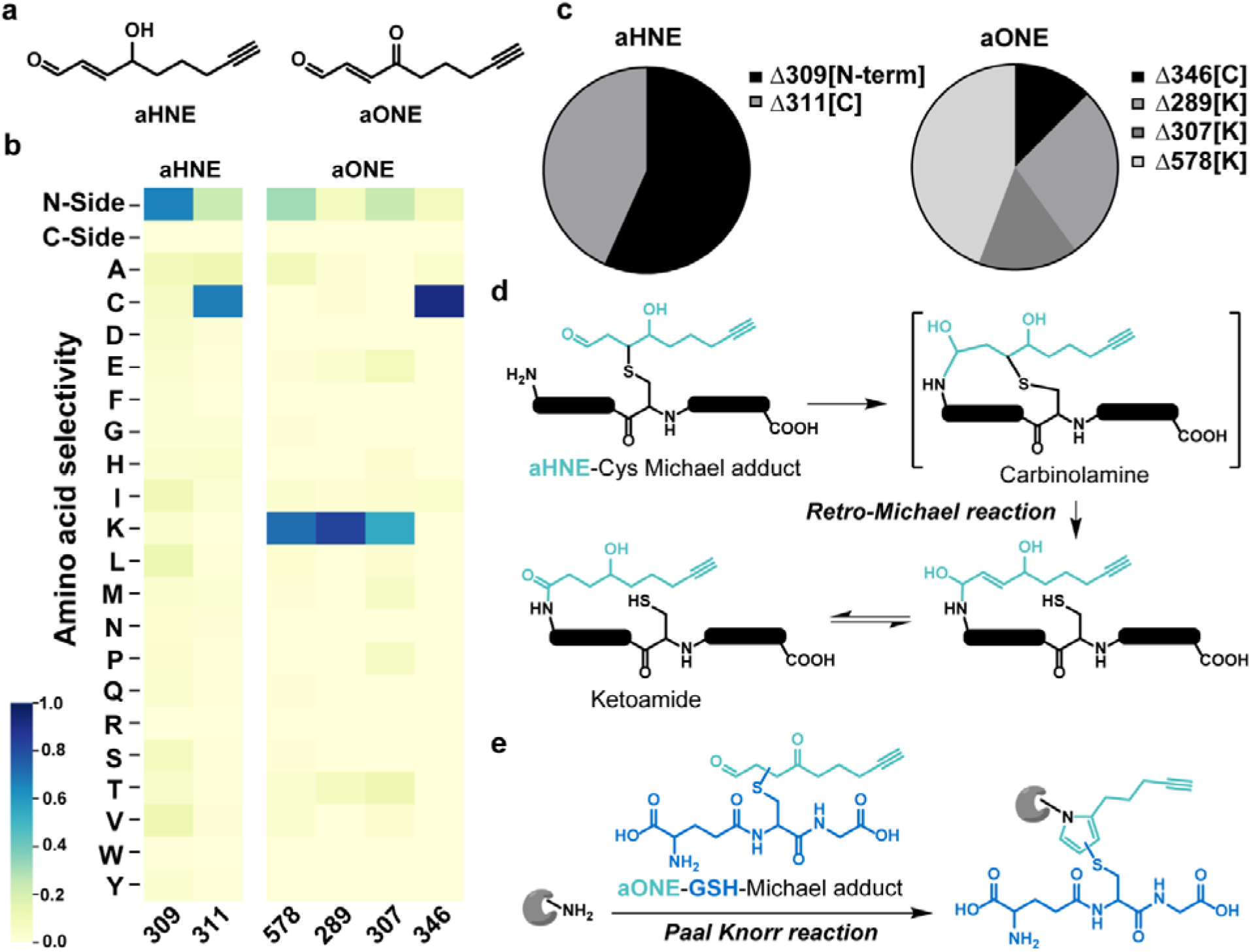
Application of pChem for lipid electrophile-derived probes. **a**, Chemical structures of the lipid electrophile-derived probes, including aHNE and aONE. **b**, Representative heatmaps showing the amino acid localization distribution of the pChem-defined PDMs for each indicated probe. **c**, Pie charts showing the abundance distribution (i.e., number of PSMs) of PDMs for each indicated probe. **d**, Plausible mechanism for the formation of an unexpected N-term modification (Δ309.17) derived from aHNE. **e**, Plausible mechanism for the formation of GSH-aONE-lysine modification (Δ578.22).

A PDM of Δ309.17 from aHNE was pinpointed by pChem on peptide N-terminal (**Fig. 5b** and **Supplementary Fig. 12**). Of interest, almost all those peptides bearing this unusual PDM contain one unmodified cysteine residue, indicative of intramolecular rearrangement (**Fig. 5d**). Adduction of aHNE to cysteinyl thiol leaves a reactive aldehyde to further react with the amino group on the corresponding peptide N-terminal, affording the carbinolamine intermediate. Followed by the retro-Michael reaction, a stable ketoamide product is formed. Hence, the cysteines on those N-term ketoamide peptide adducts could also be assigned as aHNE-modified sites. As such, ~92.0% of such cysteines were the same as those bearing the targeted PDM of Δ311.18 (Michael addition adduct).

A PDM of Δ578.22 from aONE predominantly occurred on lysine (**Fig. 5b** and **Supplementary Fig. 13**). Such a mass shift and that of aONE-derived ketoamide adduct differ by 289.08 Da, which happens to be one molecule of glutathione (GSH, a highly abundant intracellular antioxidant^56^) with a loss of water. Note that a previous report demonstrates that GSH can be rapidly cross-linked to lysines on recombinant proteins by ONE via GSH-ONE-Michael addition and subsequent Paal-Knorr condensation^57^. Likewise, the PDM of Δ578.22 should also be a *C*-glutathionylated pyrrole adduct formed in cells by the same mechanism (**Fig. 5e**). Moreover, the lysine-specificity of this PDM was also confirmed by the characteristic loss of aziridinone-based GSH-ONE-lysine conjugate (706.31 Da, **Supplementary Fig. 13**).

Taken together, the two PDMs newly uncovered by pChem allowed us to expand the aHNE/aONE-derived adductomes and to quantify their dynamics (**Supplementary Fig. 14**, **Supplementary Table 7** and **Supplementary Note 2**).

## Discussion

Unlike many other blind search engines, pChem, as a fit-for-purpose tool for chemoproteomics, is designed to produce modification-centric, constructive and easy-to-interpret outputs for probe developers.

At a cost, pChem relies on isotopic coding of PDMs that can be achieved by many well-established approaches with detailed protocols in the field of chemoproteomics (**Supplementary Fig. 1a**). For instance, many generalized isotope-coding agents, such as light and heavy clickable biotin/desthiobiotin tags are either commercially available or easy-to-synthesize (**Supplementary Fig. 1b**) ^55, 58–61^. Alternatively, a more direct way, albeit not very cost-effective, is to use isotopically labeled probes^42, 60, 62^. In addition, pChem is an .exe program free of installation, so its setup-and-run can be easily managed by probe developers, even those with no experience in informatics.

Our benchmarking results establish pChem as a robust tool by demonstrating its compatibility with a variety of data sources and probes. The analyses require neither prior knowledge nor manual inspection, thereby offering a truly unbiased survey of any modifications derived from a tested probe in complex biological systems. This unique feature would greatly accelerate the assessment of proteome-wide selectivity of chemoproteomic probes, the major rate-limiting process during their developments. Considering the continuing needs to develop residue-selective chemistry for probing protein functions^63^, discovering targeted covalent inhibitors^64^, and advancing single-molecule protein sequencing and fingerprinting technologies^65^, we consider pChem as an attractive tool for broad utilization in the field of proteomics, chemical biology and drug industry.

Moreover, together with a few previous reports^24^, the pChem analyses reported herein further strengthen the notion that endogenous reactive metabolites (e.g., formaldehyde, MGO and GSH) may possess cross-reactivity with either a probe itself or its PDMs, which has been largely overlooked in the field. This notion necessitates the use of such endogenous small molecules to benchmark chemoselectivity of a probe being developed at early stages. On the other hand, this notion would also inspire exciting pursuits to uncover previously unknown PTMs in biological systems.

Projecting forward, potential improvements of pChem include implementing it into the <u>pFind</u> studio and/or a website server, supporting data generated from alternative fragmentation techniques (e.g., electron-transfer dissociation, ETD) that can increase modification-specific fragment ions for highly labile PDMs, assessing the proteome-wide reactivity of isotopically coded cross-linkers, and so on.

## Supporting information

Supplementary Protocol

## Acknowledgements

We thank Dr. Shuai-Xin Gao and Prof. Catherine C L Wong from Peking University for their help on the use of timsTOF; Prof. Zheng-Qiu Li from Jinan University for providing the alkyl diazirine probe and for performing beta-testing; Dr. Ke Qin, Prof. Xing Chen from Peking University for providing the Ac_4_ManNAz-based glycoproteomic dataset; Mr. Peng-Yun Gong and Prof. Chao Liu for their help on the quantitative analyses of aHNE/aONE datasets; Prof. Hong-Fang Jin from Peking University for providing rat CRL-1444 cells; Prof. Xiang Xiao from Shanghai Jiaotong University for providing *E.coli* strain MG1655; Dr. Guo-Zhi Bi from Institute of Genetics and Developmental Biology, CAS, for his help in preparation of *Arabidopsis* protoplasts; Prof. Chu Wang and Prof. Xiao-Guang Lei from Peking University, Prof. Gang Li from Shenzhen Bay lab, Prof. Hui Ye from China Pharmaceutical University, Prof. Yao-Yang Zhang from Shanghai Institute of Organic Chemistry, CAS, Dr. Nan Chen from the Chomi Biotech and Dr. Uthpala Seneviratne from Pfizer for performing beta-besting. Prof. Chun-Rong Liu from Central China Normal University, Dr. Yun-Long Shi and Prof. Kate S. Carroll from the Scripps Research Institute, Prof. Peng Zou from Peking University, Prof. Qi Zhang from Fudan University, Prof. Si-Min He from Institute of Computing Technology, CAS, Prof. Yan Fu from Academy of Mathematics and Systems Science, CAS, for many insightful discussions and/or for proof-reading the manuscript. The work was supported by grants from the Natural Science Foundation of China (21922702, 81973279, and 31770885) to J.Y., (32022046) to H.C., and (32088101) to F.C.H., the National Key R&D Program of China (2016YFA0501303) to J.Y., and (2016YFA0501301) to H.C., and the State Key Laboratory of Proteomics (SKLP-K201703 and SKLP-K201804) to J.Y.

## Author contributions

J.X.H. performed the experiments, analyzed the data, and wrote the protocol, Z.C.F. designed and implemented pChem, analyze the data, and wrote the protocol, L.F., performed the QTRP experiments for various species, C.P.T. generated the AP and Diazir data sets, F.C.H. acquired funding, H.C. supervised the work, advised on pChem design and revised the manuscript. J.Y. conceived the project, supervised the work, advised on pChem design, analyzed data and the wrote the manuscript with inputs from others.

## Methods

### pChem algorithm

pChem is an installation-free .exe program relying upon an efficient database search engine termed Open-pFind^15^. It is freely available at http://pfind.ict.ac.cn/software/pChem/index.html. It can work in any desktops or laptops running Windows operating system by configuring parameters in a text editor. It can directly import the RAW data from Orbitrap instruments or read mzML files transformed by MSconvertGUI^28^ (part of ProteoWizard v.3.0.21193). Note that pChem requires DDA (data dependent acquisition)-based MS data with both MS1 and MS/MS spectra recorded in the high-resolution mode. The output files of pChem include (**Supplementary Protocol**): (1) an editable summary file containing many key characteristics of the isotopically paired PDMs, such as the number of PSMs and peptides, accurate masses, site-specific localization probability and DFLs; (2) a heat map showing the amino acid localization distribution for each PDM; (3) a radar plot showing the overall performance with the metrics of profiling efficiency, modification homogeneity and residue selectivity.

#### Blind search

pChem first automatically invokes Open-pFind^15^ to perform *bona fide* blind search against the sequence database that is provided as input. This step aims to generate abundant modification candidates at the PSM level using a tag-index strategy by which the sequence database (The FASTA canonical protein sequence databases from various species were obtained from <u>Uniprot</u>^26^) is re-constructed into the *tag-indexed protein database* and peptide candidates are retrieved. Unlike the default search mode of Open-pFind^15^, pChem does not retrieve preset modification lists (i.e., the Unimod database). For each high-resolution MS/MS spectrum, a number of *k*-mer tags are extracted with a depth-first search, which is similar to *de novo* peptide sequencing^66^. Note that the tag candidates allow modification incorporated from the pre-defined common modifications (e.g., oxidation of methionines and carbamidomethylation of cysteines). Alternatively, the MS/MS data can be initially searched by Open-pFind^15^ against Unimod to find the highly abundant non-probe derived modifications, which will then be specified and retrieved in the blind search step.

The tag candidates are then scored and only a few top-ranked ones are retained before searching against the *tag-indexed protein database*. Afterward, peptide candidates are generated by extending each of the matched tags to a full-length peptide sequence. During the extension procedure, all mass shifts within the given range (± 1,000 Da by default) are retained. Peptides that fit at least one flanking mass of each tag are further retrieved and scored at PSM-level via the same scoring strategy of Open-pFind^15^, and the non-zero mass shift (if any) on one side is considered as a potential modification. Then the mass shift is tested in turn on all sites of the peptide except those within the region matched with the tag, to generate different modified peptides. Finally, the modified peptides are scored with the matching spectra via a number of procedures (*e.g.*, reranking, FDR control and protein inference) the same as those of Open-pFind^15^.

#### Calibrating accurate masses of modification candidates

Mass shifts found by blind search are kept to only two decimal places, enabling relatively high computing efficiency. Further, a mass error calibration algorithm is implemented to obtain the highly accurate modification masses. Specifically, the *system error* of precursor ion mass is first calculated by averaging the mass difference error between the measured mass of precursor ion and its matching peptide free of unknown modification as follows:

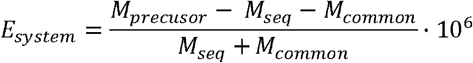

where *E*_*system*_ denotes the system error in p.p.m., *M*_*precusor*_ is the neutral mass of the precursor ion, *M*_*seq*_ is the mass of the corresponding peptide sequence and *M*_*common*_ is the summed mass of the common modifications of this peptide if exists.

For each mass shift being assigned to unknown modification at the PSM level, its accurate value can be inferenced according to the precursor ion mass calibrated with the aforementioned *system error* and the corresponding peptide sequence as follows:

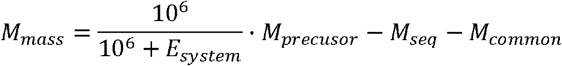

where *M*_*mass*_ denotes the accurate mass of unknown modification.

Finally, the accurate mass corresponding to the same modification candidate is unified by averaging the same calculated mass shifts from multiple PSMs as follows:

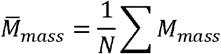

where *N* denotes the total number of PSMs containing the same modification candidate and 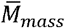 is the accurate mass computed for the modification candidate. As shown in **Supplementary Fig. 15**, pChem searches can achieve mass accuracy in the low p.p.m. range (i.e., from 0.02 to 19.7 p.p.m., with medium values of 2.4 and 3.8 p.p.m. for light and heavy PDMs characterized in this study, respectively).

#### Defining PDMs

In general, aforementioned steps will generate and output numerous modification candidates (typically from dozens to several hundreds, **Supplementary Fig. 16**), thereby complicating the result interpretation. To this end, several semi-empirical criteria are applied to automatically recognize the genuine PDMs. First, the isotope coding information is utilized to eliminate non-PDMs or other unrealistic modifications. For the six-heavy carbon coding strategy, the theoretical mass difference between a pair of light and heavy PDMs is 6.020132 Da. If such a measured mass difference is out of the range of [6.020132 − 0.001, 6.020132 + 0.001] Da, the modification will be neglected. Second, less abundant modification candidates with the PSM counting number lower than a predefined threshold (*i.e.*, 5% of total, by default. As a result, the PSMs of the *high-confidence PDMs* generally account for >85% of those of all *isotope-paired PDMs*, **Supplementary Fig. 17**) are also filtered out. Third, only the modifications with masses larger than the pre-defined threshold, *e.g.*, 200 Da by default, are retained. By applying such criteria, typically, less than five high-confidence PDM pairs will be narrowed down and reported in the final output summary. For all probes tested herein, these high-confidence PDMs account for only 2.3% (median, ranging from 0.63% to 13.8%) of initial modification candidates by open-search (**Supplementary Fig. 16**), thereby facilitating result interpretation by users.

#### Localizing PDMs

Once PDMs are defined, the subsequent statistical analysis is carried out to estimate the corresponding localization probability by calculating the position distribution for each PDM as follows:

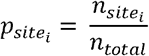

In this formula, 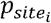 denotes the localization probability of PDM occurring at *site*_*i*_, 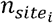 is the number of PSMs related to each PDM that occur at specific *site*_*i*_, and *n*_*total*_ denotes the total number of PSMs related to the same PDM. The positions of the top-ranked 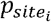 are reported in *pChem.summary* file. Meanwhile, for each PDM, a heat map showing 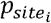 values on all possible sites will be generated automatically.

Note, in addition to twenty protein-coding amino acids, peptide N- and C-termini are also considered in the modification site determination. Specifically, given a modified peptide whose length is *L*, if modification located at the leftmost amino acid (position 1), it is also regarded as a potential N-terminal modification with a position identifier of 0. Similarly, if the modification located at the rightmost amino acid (position *L*), it is also regarded as a potential C-terminal modification with a position identifier of *L* + 1. For example, given a peptide sequence CEHVAEADK, if a PDM is identified on the first amino acid C with blind search, the modification is also considered to be located at the N-terminal of the peptide by the algorithm.

#### Recognizing diagnostic fragment losses (DFLs)

Without prior knowledge of the fragmentation patterns, an unsupervised outlier-detection algorithm^66^ can be adopted to automatically recognize potential PDM-specific DFL set (*Δ*= {*δ*_1_, … , *δ*_*k*_}) by searching a collection of MS/MS spectra corresponding to each isotopic pair of PDMs. First, for each PSM related to a PDM, all theoretical *b*- and *y*-ion masses *T* = {*t*_1_, … , *t*_*n*_} are calculated. For example, given a peptide VQ<u>S</u>VEK and the PDM is identified on S, the masses of the theoretical ions with the PDM is referred to as *T*′ = {*b*_3_~*b*_5_, *y*_4_~*y*_5_}. Let *S*= {*s*_1_, … , *s*_*m*_} be the experimental spectrum peak set. A theoretical fragment ion mass *t*_*i*_ and an experimental peak *s*_*j*_ have an offset *x*_*i,j*_ = *t*_*i*_ − *s*_*j*_, which can be considered as a discrete random variable. Since the frequencies of the offsets corresponding to the DFLs are expected to be much larger than those of the random offsets, the peaks in the empirical distribution of the offsets allow to reveal the genuine PDM-specific DFLs. The statistics of offsets over all experimental spectrum peaks and all theoretical peaks provides a reliable learning algorithm to generate DFL set Δ, whose frequency should exceed the half of top one. That is, if the isotopically paired PDMs also generate DFLs that can be paired as mentioned above and the unmatched DFLs will be filtered out as noises (**Supplementary Fig. 18a**). Instead, if the offset with the highest frequency is equal to zero, it suggests that the corresponding PDM is unlikely to generate any significant DFLs (**Supplementary Fig. 18b**). Note that DFL set Δ can be further refined by applying the isotopic mass difference cutoff value as above.

#### Restricted search

The characteristics of the PDMs (*i.e.*, the accurate mass and the Top1 localization site) are automatically retrieved by pChem for the last round of restricted search in pFind 3, so are the predefined common modifications. This procedure further increases the spectrum identification rate that is measured by the percentage of spectra identified by the search engine among all input spectra within the threshold of 1% FDR estimated by the target-decoy strategy^67^ at the peptide level, thereby facilitating the performance scoring process (see below). Moreover, the restricted search can revise the identification when a PDM and one or more adjacent common modifications on the same peptide sequence are erroneously identified as a ‘new’ PDM by summing up masses of them.

#### Probe performance scoring

For tool users such as probe developers, pChem provides a general evaluation of the performance of the tested probe by scoring values as follows:

1. *Profiling efficiency (%)*, which indicates the capability of the PDMs being identified from the MS data and is calculated as:

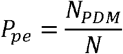

where *N*_*PDM*_ represents the number of all PSMs assigned to the isotopically paired PDMs (limited to top-5 PDMs by default) and *N* is the total number of PSMs identified in the restricted search procedure.
2. *Modification homogeneity (%),* which measures the modification location uniformity of a PDM and is computed as:

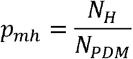

where *N*_*H*_ denotes the number of the PSMs assigned to the isotopically paired PDM with the most identified PSMs.
3. *Residue selectivity (%),* which evaluates the position selectivity attribution of PDM and can be formed as:

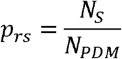

where *N*_*S*_ corresponds to the number of PSMs corresponding to the site with the highest number of identified PSMs assigned to the PDMs. Note that the latter two scores are calculated right after the blind search procedure.

### Chemoproteomic data sets published previously

Raw data files previously generated by using the IPM-based QTRP method are available from ProteomeXchange (PXD027762 for evaluating quantification accuracy^4^; PXD027767 for profiling the cysteinome in *Drosophila melanogaster*^68^, *Caenorhabditis elegans*^43^, *Psedomonas syringae* ^69^, and *Mus musculus* ^70^). Raw analyses of chemoproteomic data sets based on oxoform-specific probes (i.e., DYn-2, BTD, WYnes and DiaAlk) have been described previously and are available now from ProteomeXchange (PXD027764)^12–13, 42, 49–50^. The aHNE- and aONE-based chemoproteomic analyses have also been reported previously^24, 55^, and the corresponding raw data sets can be obtained from ProteomeXchange (PXD027760 and PXD007149, respectively). Raw data files for the Ac_4_ManNAz-based glycoproteomic analysis were generated and kindly provided by Prof. Xing Chen and colleagues^59^.

### Chemoproteomic data sets newly generated in this study

Raw data files newly generated by using the IPM-based QTRP method are available from ProteomeXchange (PXD027755 for initial test; PXD027758 for data collected from different instruments; PXD027767 for profiling the cysteinome in *Arabidopsis thaliana*, *Escherichia coli*, and *Rattus norvegicus*). QTRP data files generated by using other thiol-reactive probes (i.e., ENE, NPM, PPMS, VSF, and ‘cocktail’) are available from ProteomeXchange (PXD027756). Raw data files from the NHS- and STP-based lysinome profiling are available from ProteomeXchange (PXD027789). The AP- and Diazir-based chemoproteomic data sets are available from ProteomeXchange (PXD027591).

### Sample preparation

All probes used here contain a ‘clickable’ alkyne tag, one of most-used functionalized handles for chemoproteomics^1^. For pChem search, a well-established quantitative chemoproteomic workflow relying on the commercially available light and heavy azido-biotin reagents with a photocleavable linker was utilized to isotopically code alkyne probe-derived modifications (**Supplementary Fig. 1b**). More details are described as follows.

#### Reagents

IPM, ENE and VSF were house-made as previously described^71–72^. NPM (cat. No. TA113) was purchased from KeraFast. PPMS (cat. No. P757300) was purchased from Toronto research chemicals. STP (cat. No. 30720) was purchased from Lumiprobe. NHS (cat. No. A171448) was purchased from Aladdin. AP (9186096) was purchased from J&K Scientific. The Diazir probe was kindly provided by Prof. Zheng-Qiu Li^73^. Iodoacetamide (IAA, cat. No. V900335), tris[(1-benzyl-1H-1,2,3-triazol-4-yl)methyl]amine (TBTA) (cat. No. 678937), and sodium ascorbate (cat. No. A7631) were purchased from Sigma-Aldrich.

Dithiothreitol (DTT, cat. No. A620058-0025) was purchased from BBI Life Sciences; Light Azido-UV-Biotin (cat. No. EVU102), and Heavy Azido-UV-Biotin (cat. No. EVU151) were purchased from KeraFast; Sequencing grade trypsin (cat. No. V5113) was purchased from Promega. Streptavidin sepharose high performance (cat. No. 17-5113-01) was purchased from GE. All other reagents were purchased from Thermo Fisher Scientific unless otherwise noted.

#### Preparation of cell lysates

Human HEK293T cells and rat CRL-1444 cells were purchased from the National Infrastructure of Cell Line Resource (China), cultured in DMEM supplemented with 10% FBS (cat. No. 10099141), 1% penicillin-streptomycin (Cell World, cat. No. C0160-611) and 1% Glutagro (Corning, cat. No. 25-015-CI), maintained at 37°C in a 5% CO_2_ humidified atmosphere. Cells were grown to ~80% confluency, washed with prechilled PBS, and lysed in pre-chilled lysis buffer (50 mM HEPES (pH 7.6), 150 mM NaCl, and 1% IGEPAL) supplemented with 1 x protease and phosphatase inhibitors (cat. No. A32961).

*E.coli* strain MG1655 was cultivated 50 mL M9 minimal media containing 0.2% casamino acids and 10 mM glucose to an optical density at 600 nm of ~0.3. 4 mL of culture was then transferred into 15 mL conical centrifuge tubes with 5× minimum inhibitory concentration (MIC) of antibiotics and incubated for additional 4h. The culture was centrifuged at 1,400 × g for 10 min. Cell pellets were harvested by centrifugation and lysed as above.

*Arabidopsis* protoplasts were prepared as previously described^74^. In brief, well-expanded leaves from *Arabidopsis thaliana* (3-4 weeks) were sliced into small leaf strips (0.5-1 mm) and incubated in 20 mM MES (pH 5.7), 1.5% (w/v) cellulase R10, 0.4% (w/v), macerozyme R10, 0.4 M mannitol, 20 mM KCl, 10 mM CaCl_2_, and 0.1% BSA. The leaf strips were vacuum-infiltrated in a desiccator for 30 min and incubated for additional 3h at RT in the dark. The digested sample was then filtrated with a 75 µm nylon mesh. The protoplasts were pelleted by centrifugation at 100g for 2 min, resuspended in W5 solution (2 mM MES, pH 5.7, 154 mM NaCl, 125 mM CaCl_2_ and 5 mM KCl), pelleted again on ice for 30 min by gravity, and lysed in prechilled HEPES buffer (50 mM, pH 7.6) containing 150 mM NaCl, 1mM EDTA, 0.5% Triton X-100, and 1% IGEPA, 1 x protease and phosphatase inhibitors.

#### Preparation of the probe-labeled proteomes

The cell lysates were then incubated with each probe as indicated with the conditions (i.e., concentration, time, temperature) summarized in **Supplementary Table 8**. The resulting protein samples were reduced with DTT (10 mM, 1 h, RT), and subsequently alkylated with IAA (40 mM, 1 h, RT, with light protection). Proteins were then precipitated with a methanol-chloroform system (aqueous phase/methanol/chloroform, 4:4:1 (v/v/v)). With sonication, the precipitated proteins were resuspended in 50 mM ammonium bicarbonate, and digested with trypsin at a 1:50 (enzyme/substrate) ratio overnight at 37°C. The tryptic digests were desalted with HLB extraction cartridges (Waters, cat. No. 186000383), dried under vacuum, and resuspended in a water solution containing 30% acetonitrile. CuAAC reaction was performed at RT for 2 h with rotation and light protection by subsequently adding 1 mM either light or heavy Azido-UV-biotin (1 μL of a 40 mM stock), 10 mM sodium ascorbate (4 μL of a 100 mM stock), 1 mM TBTA (1 μL of a 50 mM stock), and 10 mM CuSO_4_ (4 μL of a 100 mM stock). The light and heavy isotopic tagged samples were then mixed immediately following CuAAC, cleaned with strong cation exchange (SCX, Nest group, cat. No SMM HIL-SCX) spin columns, and then enriched with streptavidin for 2 h at RT. Streptavidin beads were then washed with 50 mM NaAc (pH=4.5), 50 mM NaAc containing 2 M NaCl (pH=4.5), and deionized water twice each with end-to-end rotations, then resuspended in 25 mM ammonium bicarbonate, transferred to glass tubes (VWR), and irradiated with UV lamp at 365 nm (2 h, RT, with magnetic stirring). The supernatant was collected, concentrated under vacuum, and desalted with HLB cartridges. The resulting peptides were evaporated to dryness and reconstituted in 0.1% formic acid for LC-MS/MS analysis.

### LC-MS/MS

The LC-MS/MS data sets (incl. 44 representative files) used here for benchmarking pChem were generated on ten different LC-MS/MS instruments by three vendors from four independent laboratories (**Supplementary Table 9**). The settings for each LC-MS/MS instrument are summarized in **Supplementary Table 10**.

### Data analysis

For pChem search, the running time depends on the computer configuration, the complexity of the tested probe-label peptide samples, and the types and settings of LC-MS/MS instruments. Here, all analyses were performed on a desktop PC running Microsoft Window 10 Home (v.19041.110), with one 2.90GHz CPU Intel i7-10700 processor with 64 GB of installed RAM. In this computing system, for example, approximately 20 min would be required for the default pChem search of the data generated from a single 75-min EasyLC tandem Q-Exactive plus analysis of the IPM-based QTRP sample. The running times used for pChem search of MS data generated from other instruments are shown in **Supplementary Fig. 3a**.

For the pFind 3-based re-analyses of aHNE and aONE data sets, MS data files were searched against *Homo sapiens* Uniprot canonical database (Downloaded on May 16, 2021, 20,395 entries). Precursor ion mass and fragmentation tolerance were set as ±10 p.p.m. and ±20 p.p.m., respectively. The maximum number of modifications allowed per peptide was three, as was the maximum number of missed cleavages allowed. Common modifications (i.e., oxidation of methionines and carbamidomethylation of cysteines) and PDMs (refer to **Supplementary Table 1**) were all searched as variable modifications. A differential modification of 6.020132 Da on PDM was used for stable-isotopic quantification. The FDRs at spectrum, peptide, and protein level were ≤1%. The MS1-based quantification was performed using pQuant^75^, which calculated R_H/L_ values based on each identified MS scan with a 15 p.p.m.-level m/z tolerance window and assigned an interference score (σ) to each value from zero to one. In principle, the lower the calculated σ was, the less co-elution interference signal was observed in the extracted ion chromatograms. In this regard, the median values of probe-modified peptide ratios with σ less than or equal 0.5 were considered to calculate site-level ratios. Quantification results were obtained from three biological replicates with two LC-MS/MS runs for each.

## Data Availability

The newly generated chemoproteomic data sets have been deposited to the ProteomeXchange Consortium via the PRIDE^76^ partner repository with the dataset identifiers PXD027755 (Username, Password: mub4eIda), PXD027758 (Username, Password: 46z1w0cj), PXD027767 (Username, Password: EZL1H061), PXD027789 (Username, Password: T9s2rlJ3), PXD027756 (Username, Password: IeSLiwOa); previously published data were also used to benchmark pChem in repositories with identifiers PXD027591 (Username, Password: WgBqKEGi; PXD007149), PXD027764 (Username, Password: Zje4S2sN), PXD027762 (Username, Password: vLSe4BNQ), PXD027760 (Username, Password: djuIY94L)

## Code availability

pChem is open-source and is freely available at https://github.com/pFindStudio/pChem under a permissive license.

## References

1. Parker, C.G. & Pratt, M.R. Click Chemistry in Proteomic Investigations. Cell 180, 605–632 (2020).

2. Grammel, M. & Hang, H.C. Chemical reporters for biological discovery. Nat Chem Biol 9, 475–484 (2013).

3. Weerapana, E. et al. Quantitative reactivity profiling predicts functional cysteines in proteomes. Nature 468, 790–795 (2010).

4. Fu, L. et al. A quantitative thiol reactivity profiling platform to analyze redox and electrophile reactive cysteine proteomes. Nat Protoc 15, 2891–2919 (2020).

5. Hacker, S.M. et al. Global profiling of lysine reactivity and ligandability in the human proteome. Nat Chem 9, 1181–1190 (2017).

6. Hahm, H.S. et al. Global targeting of functional tyrosines using sulfur-triazole exchange chemistry. Nat Chem Biol 16, 150–159 (2020).

7. Lin, S. et al. Redox-based reagents for chemoselective methionine bioconjugation. Science 355, 597–602 (2017).

8. Taylor, M.T., Nelson, J.E., Suero, M.G. & Gaunt, M.J. A protein functionalization platform based on selective reactions at methionine residues. Nature 562, 563–568 (2018).

9. Jia, S., He, D. & Chang, C.J. Bioinspired Thiophosphorodichloridate Reagents for Chemoselective Histidine Bioconjugation. J Am Chem Soc 141, 7294–7301 (2019).

10. Ma, N. et al. 2H-Azirine-Based Reagents for Chemoselective Bioconjugation at Carboxyl Residues Inside Live Cells. J Am Chem Soc 142, 6051–6059 (2020).

11. Bach, K., Beerkens, B.L.H., Zanon, P.R.A. & Hacker, S.M. Light-Activatable, 2,5-Disubstituted Tetrazoles for the Proteome-wide Profiling of Aspartates and Glutamates in Living Bacteria. ACS Cent Sci 6, 546–554 (2020).

12. Akter, S. et al. Chemical proteomics reveals new targets of cysteine sulfinic acid reductase. Nat Chem Biol 14, 995–1004 (2018).

13. Gupta, V., Yang, J., Liebler, D.C. & Carroll, K.S. Diverse Redoxome Reactivity Profiles of Carbon Nucleophiles. J Am Chem Soc 139, 5588–5595 (2017).

14. Tian, C., Liu, K., Sun, R., Fu, L. & Yang, J. Chemoproteomics Reveals Unexpected Lysine/Arginine-Specific Cleavage of Peptide Chains as a Potential Protein Degradation Machinery. Anal Chem 90, 794–800 (2018).

15. Chi, H. et al. Comprehensive identification of peptides in tandem mass spectra using an efficient open search engine. Nat Biotechnol 36, 1059–1061 (2018).

16. Devabhaktuni, A. et al. TagGraph reveals vast protein modification landscapes from large tandem mass spectrometry datasets. Nat Biotechnol 37, 469–479 (2019).

17. Geiszler, D.J. et al. PTM-Shepherd: Analysis and Summarization of Post-Translational and Chemical Modifications From Open Search Results. Mol Cell Proteomics 20, 100018 (2020).

18. Kong, A.T., Leprevost, F.V., Avtonomov, D.M., Mellacheruvu, D. & Nesvizhskii, A.I. MSFragger: ultrafast and comprehensive peptide identification in mass spectrometry-based proteomics. Nat Methods 14, 513–520 (2017).

19. Chick, J.M. et al. A mass-tolerant database search identifies a large proportion of unassigned spectra in shotgun proteomics as modified peptides. Nat Biotechnol 33, 743–749 (2015).

20. Yu, F. et al. Identification of modified peptides using localization-aware open search. Nat Commun 11, 4065 (2020).

21. Yu, F., Li, N. & Yu, W. PIPI: PTM-Invariant Peptide Identification Using Coding Method. J Proteome Res 15, 4423–4435 (2016).

22. Na, S., Bandeira, N. & Paek, E. Fast multi-blind modification search through tandem mass spectrometry. Mol Cell Proteomics 11, M111 010199 (2012).

23. Dasari, S. et al. TagRecon: high-throughput mutation identification through sequence tagging. J Proteome Res 9, 1716–1726 (2010).

24. Sun, R. et al. Chemoproteomics Reveals Chemical Diversity and Dynamics of 4-Oxo-2-nonenal Modifications in Cells. Mol Cell Proteomics 16, 1789–1800 (2017).

25. Zanon, P.R.A. et al. Profiling the proteome-wide selectivity of diverse electrophiles ChemRxiv Preprint at 10.26434/chemrxiv.14186561.v1 (2021).

26. Creasy, D.M. & Cottrell, J.S. Unimod: Protein modifications for mass spectrometry. Proteomics 4, 1534–1536 (2004).

27. Reisz, J.A., Bechtold, E., King, S.B., Poole, L.B. & Furdui, C.M. Thiol-blocking electrophiles interfere with labeling and detection of protein sulfenic acids. FEBS J 280, 6150–6161 (2013).

28. Chambers, M.C. et al. A cross-platform toolkit for mass spectrometry and proteomics. Nat Biotechnol 30, 918–920 (2012).

29. Wu, C.W. & Yarbrough, L.R. N-(1-pyrene)maleimide: a fluorescent cross-linking reagent. Biochemistry 15, 2863–2868 (1976).

30. Halloran, M.W. & Lumb, J.P. Recent Applications of Diazirines in Chemical Proteomics. Chemistry 25, 4885–4898 (2019).

31. West, A.V. et al. Labeling Preferences of Diazirines with Protein Biomolecules. J Am Chem Soc 143, 6691–6700 (2021).

32. Qin, W., Cho, K.F., Cavanagh, P.E. & Ting, A.Y. Deciphering molecular interactions by proximity labeling. Nat Methods 18, 133–143 (2021).

33. Bar, D.Z. et al. Biotinylation by antibody recognition-a method for proximity labeling. Nat Methods 15, 127–133 (2018).

34. Udeshi, N.D. et al. Antibodies to biotin enable large-scale detection of biotinylation sites on proteins. Nat Methods 14, 1167–1170 (2017).

35. Li, Y. et al. A Clickable APEX Probe for Proximity-Dependent Proteomic Profiling in Yeast. Cell Chem Biol 27, 858–865 e858 (2020).

36. Massari, J. et al. Acetyl radical production by the methylglyoxal-peroxynitrite system: a possible route for L-lysine acetylation. Chem Res Toxicol 23, 1762–1770 (2010).

37. Tokikawa, R., Loffredo, C., Uemi, M., Machini, M.T. & Bechara, E.J. Radical acylation of L-lysine derivatives and L-lysine-containing peptides by peroxynitrite-treated diacetyl and methylglyoxal. Free Radic Res 48, 357–370 (2014).

38. Ross, D. & Moldeus, P. Generation of reactive species and fate of thiols during peroxidase-catalyzed metabolic activation of aromatic amines and phenols. Environ Health Perspect 64, 253–257 (1985).

39. Sadler, A., Subrahmanyam, V.V. & Ross, D. Oxidation of catechol by horseradish peroxidase and human leukocyte peroxidase: reactions of o-benzoquinone and o-benzosemiquinone. Toxicol Appl Pharmacol 93, 62–71 (1988).

40. Paulsen, C.E. & Carroll, K.S. Cysteine-mediated redox signaling: chemistry, biology, and tools for discovery. Chem Rev 113, 4633–4679 (2013).

41. Alcock, L.J., Perkins, M.V. & Chalker, J.M. Chemical methods for mapping cysteine oxidation. Chem Soc Rev 47, 231–268 (2018).

42. Yang, J., Gupta, V., Carroll, K.S. & Liebler, D.C. Site-specific mapping and quantification of protein S-sulphenylation in cells. Nat Commun 5, 4776 (2014).

43. Meng, J. et al. Global profiling of distinct cysteine redox forms reveals wide-ranging redox regulation in C. elegans. Nat Commun 12, 1415 (2021).

44. Huang, J. et al. Mining for protein S-sulfenylation in Arabidopsis uncovers redox-sensitive sites. Proc Natl Acad Sci U S A 116, 21256–21261 (2019).

45. Huang, Y. et al. Endogenous SO2-dependent Smad3 redox modification controls vascular remodeling. Redox Biol 41, 101898 (2021).

46. Pople, J.M.M. & Chalker, J.M. A critical evaluation of probes for cysteine sulfenic acid. Curr Opin Chem Biol 60, 55–65 (2021).

47. Shi, Y. & Carroll, K.S. Comments on ‘A critical evaluation of probes for cysteine sulfenic acid’. Curr Opin Chem Biol 60, 131–133 (2021).

48. Yang, J. et al. Global, in situ, site-specific analysis of protein S-sulfenylation. Nat Protoc 10, 1022–1037 (2015).

49. Fu, L., Liu, K., Ferreira, R.B., Carroll, K.S. & Yang, J. Proteome-Wide Analysis of Cysteine S-Sulfenylation Using a Benzothiazine-Based Probe. Curr Protoc Protein Sci 95, e76 (2019).

50. Shi, Y., Fu, L., Yang, J. & Carroll, K.S. Wittig reagents for chemoselective sulfenic acid ligation enables global site stoichiometry analysis and redox-controlled mitochondrial targeting. Nat Chem (2021) **https://www.nature.com/articles/s41557-021-00767-2.**

51. Ramasamy, R., Yan, S.F. & Schmidt, A.M. Methylglyoxal comes of AGE. Cell 124, 258–260 (2006).

52. Palaniappan, K.K. & Bertozzi, C.R. Chemical Glycoproteomics. Chem Rev 116, 14277–14306 (2016).

53. Qin, W. et al. Artificial Cysteine S-Glycosylation Induced by Per-O-Acetylated Unnatural Monosaccharides during Metabolic Glycan Labeling. Angew Chem Int Ed Engl 57, 1817–1820 (2018).

54. Sayre, L.M., Lin, D., Yuan, Q., Zhu, X. & Tang, X. Protein adducts generated from products of lipid oxidation: focus on HNE and one. Drug Metab Rev 38, 651–675 (2006).

55. Yang, J., Tallman, K.A., Porter, N.A. & Liebler, D.C. Quantitative chemoproteomics for site-specific analysis of protein alkylation by 4-hydroxy-2-nonenal in cells. Anal Chem 87, 2535–2541 (2015).

56. Meister, A. & Anderson, M.E. Glutathione. Annu Rev Biochem 52, 711–760 (1983).

57. Zhu, X., Gallogly, M.M., Mieyal, J.J., Anderson, V.E. & Sayre, L.M. Covalent cross-linking of glutathione and carnosine to proteins by 4-oxo-2-nonenal. Chem Res Toxicol 22, 1050–1059 (2009).

58. Zanon, P.R.A., Lewald, L. & Hacker, S.M. Isotopically Labeled Desthiobiotin Azide (isoDTB) Tags Enable Global Profiling of the Bacterial Cysteinome. Angew Chem Int Ed Engl 59, 2829–2836 (2020).

59. Qin, K. et al. Quantitative Profiling of Protein O-GlcNAcylation Sites by an Isotope-Tagged Cleavable Linker. ACS Chem Biol 13, 1983–1989 (2018).

60. Abo, M., Li, C. & Weerapana, E. Isotopically-Labeled Iodoacetamide-Alkyne Probes for Quantitative Cysteine-Reactivity Profiling. Mol Pharm 15, 743–749 (2018).

61. Li, J. et al. An Isotope-Coded Photocleavable Probe for Quantitative Profiling of Protein O-GlcNAcylation. ACS Chem Biol 14, 4–10 (2019).

62. VanHecke, G.C., Yapa Abeywardana, M., Huang, B. & Ahn, Y.H. Isotopically Labeled Clickable Glutathione to Quantify Protein S-Glutathionylation. Chembiochem 21, 853–859 (2020).

63. Niphakis, M.J. & Cravatt, B.F. Enzyme inhibitor discovery by activity-based protein profiling. Annu Rev Biochem 83, 341–377 (2014).

64. Chan, W.C., Sharifzadeh, S., Buhrlage, S.J. & Marto, J.A. Chemoproteomic methods for covalent drug discovery. Chem Soc Rev (2021).

65. Alfaro, J.A. et al. The emerging landscape of single-molecule protein sequencing technologies. Nat Methods 18, 604–617 (2021).

## References

66. Dančík, V., et al. De novo peptide sequencing via tandem mass spectrometry. J Comput Biol 6, 327–342 (1999).

67. Elias, J.E., Gygi, S.P. Target-decoy search strategy for increased confidence in large-scale protein identifications by mass spectrometry. Nat Methods 4, 207–214 (2007).

68. Petrova B, et al. Dynamic redox balance directs the oocyte-to-embryo transition via developmentally controlled reactive cysteine changes. Proc Natl Acad Sci U S A 115, E7978–E7986 (2018)

69. Wang W, et al. An Arabidopsis Secondary Metabolite Directly Targets Expression of the Bacterial Type III Secretion System to Inhibit Bacterial Virulence. Cell Host Microbe 27, 601–613.e7 (2020).

70. Sun, R., et al. A Chemoproteomic Platform to Assess Bioactivation Potential of Drugs. Chem Res Toxicol 30, 1797–1803 (2017).

71. Lin, D., Saleh, S., & Liebler, D.C. Reversibility of covalent electrophile-protein adducts and chemical toxicity. Chem Res Toxicol 21, 2361–2369 (2008).

72. Weerapana, E., Simon, G.M., & Cravatt B.F. Disparate proteome reactivity profiles of carbon electrophiles. Nat Chem Biol 4, 405–407 (2008).

73. Pan, S., et al. A Suite of “Minimalist” Photo-Crosslinkers for Live-Cell Imaging and Chemical Proteomics: Case Study with BRD4 Inhibitors. Angew Chem Int Ed Engl. 56, 11816–11821 (2017)

74. Yoo, S.D., Cho, Y.H., & Sheen, J. Arabidopsis mesophyll protoplasts: a versatile cell system for transient gene expression analysis. Nat Protoc 2,1565–1572 (2007).

75. Liu, C., et al. pQuant improves quantitation by keeping out interfering signals and evaluating the accuracy of calculated ratios. Anal Chem 86, 5286–5294 (2014).

76. Perez-Riverol, Y., et al. The PRIDE database and related tools and resources in 2019: improving support for quantification data. Nucleic Acids Res 47, D442–D450 (2019)

