## Supplementary Protocol for "pChem: a modification-centric assessment tool for the performance of chemoproteomic probes"

### User Manual for pChem

**6. Supporting Protocol 2** .......... **11-13**

### 1. Download

1.1. pChem can be freely downloaded from
http://pfind.ict.ac.cn/software/pChem/index.html

#### pChem

Introduction - Cite us - Downloads

##### Introduction

Chemical probe coupled with mass spectrometry (MS)-based proteomics, herein termed chemoproteomics, offers versatile tools to globally profile protein features and to systematically interrogate the mode of action of small molecules in a native biological system. Nonetheless, development of an efficient and selective probe for chemoproteomics can still be challenging. Besides, it is also difficult to unbiasedly assess its chemoselectivity at a proteome-wide scale. Here we present pChem, a modification-centric blind search and summarization tool to provide a pipeline for rapid and unbiased assessing of the performance of ABPP and metabolic labeling probes. This pipeline starts experimentally by isotopic coding of PDMs, which can be automatically recognized, paired, and accurately reported by pChem, further allowing users to score the profiling efficiency, modification-homogeneity and proteome-wide residue selectivity of a chemoproteomic probe.

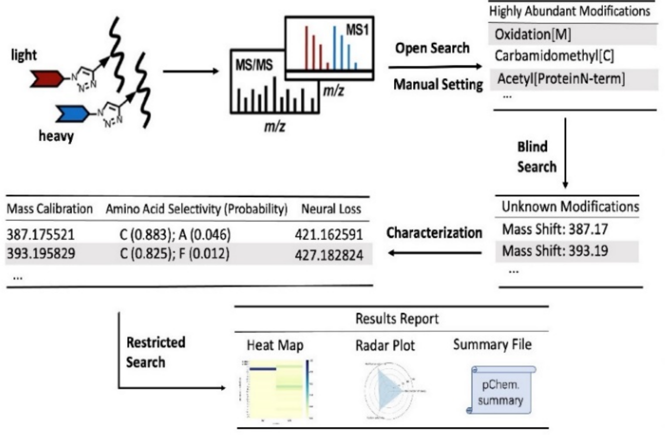

##### Cite us

pChem: a modification-centric assessment tool for performance of chemoproteomic probes.
Ji-Xiang He, Zheng-Cong Fei, Fu-Chu He, Si-Min He, Hao Chi, Jing Yang.
Under review

##### Downloads

pChem 1.0 is currently free to use. click to download.

For source code, please refer to github.

For detailed usage, please refer to user guide.

For other issues, please contact.

1.2. click "*click to download*"

1.3. Un-compress the "pChem.zip" package into a specified file folder
pChem.zip

#### 2. Configuration

2.1. Double click *pChem* 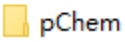

to open the folder

2.2. Open "*pChem.cfg*" using a text editor, *e.g.*, Microsoft Notepad or Notepad++ (recommend) (https://notepad-plus.en.softonic.com/)

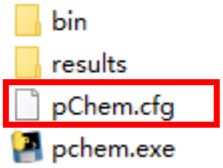

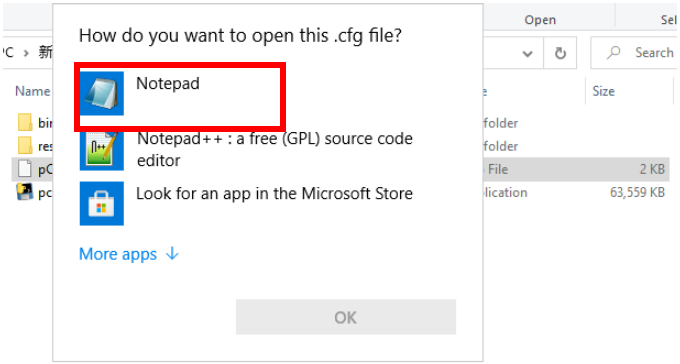

2.3. Parameter setting in "*pChem.cfg*"

① # Path to install pFind software

pfind_install_path=X:\pChem\bin\pfind

**Illustration:** the executable files of pFind are stored in pChem\bin\pfind
**Note:** The blank space needs to be avoided. The same advice holds true for ②-④.

② # Path to the output file

output_path=X:\pChem\results

**Note:** The users are suggested to change this path to output file folder when starting a new project.

③ # Path to the protein sequence database

fasta_path=X:\XXX\XXX.fasta

**Note:** Protein *.fasta database of the targeted species (e.g., *Home sapiens*) can be downloaded from Uniprot as described in **Supporting Protocol 1**.

④ # Format of MS data, RAW or MZML

msmstype=RAW

**Illustration: Default**
**Note:** Non-Thermo MS data need to be converted into mzML files before pCheml search. The users can refer to **Supporting Protocol 2**.

⑤ # The number and path of MS data

msmsnum=N

msmspath1=X:\XXX\XXX.RAW

msmspath2=X:\XXX\XXX.RAW

……………………………………

msmspathN=X:\XXX\XXX.RAW

⑥ # Type of MS dissociation method

activation_type=HCD-FTMS

**Illustration: Default**
**Note:** There is no need to change this setting if MS data is generated by TOF instruments.

⑦ # Usage of open search (True/ False), the common modification can be set if not

open_flag=False

common_modification_number=2

common_modification_list=Carbamidomethyl[C];Oxidation[M];

**Illustration: Default**
**Note:** The names of common modifications should be the same as those appeared in Unimod database.

⑧ # Mass tolerance of the mass shift between light isotope and heavy isotope

mass_of_diff_diff=6.020132

**Note:** This default value is calculated based on the isotopic mass shift between six heavy and light carbons encoded in probe-derived modifications (PDMs). Users can set any other values based on their different isotope labeling strategies. Monoisotopic masses of elements commonly used for coding PDMs are listed in the Table below.
**Troubleshooting:** One always needs to confirm this value being correctly input.

| Light | Monoisotopic mass | Heavy | Monoisotopic mass |
|---|---|---|---|
| H | 1.007825 | $^{2}H$ | 2.014102 |
| C | 12.000000 | $^{13}C$ | 13.003355 |
| N | 14.003073 | $^{15}N$ | 15.000109 |
| O | 15.994914 | $^{18}O$ | 17.999162 |

⑨ # Isotopic mass difference within empirically defined tolerance(ppm)

mass_diff_diff_range=166

**Illustration: Default**

**Note:** For 6.020132 Da of isotopic mass difference, ± 0.001 Da of mass tolerance represents 166 p.p.m. (=1000,000*0.001Da/6.020132Da).

**Troubleshooting:** If the pChem search mis-identified the targeted PDMs or even report nothing, one might want to loose the defined mass tolerance (e.g., 830 ppm for 0.005 Da of mass difference).

⑩ # Minimum mass for unknown modification (Da)

min_mass_modification=200

**Illustration: Default**

**Note:** The PDMs generated from the use of bioorthogonal cleavable linkers typically possess masses higher than 200 Da.

⑪ # Isotopic pairs of mass shifts with PSMs less than X% of that of overall PDMs were neglected

filter_frequency=5

**Illustration: Default**

**Note:** This parameter can be set as 0 if one wants to retrieve all PDMs including those with just a few PSMs.

⑫ # if consider the N- or C-termini for amino acid localization (True or false)

side_position=True

**Illustration: Default**

⑬ # if use restricted search for radar figure plotting (True or false)

use_close_search=True

**Illustration: Default**

#### 3. Run

Once all parameters have being set, double click "*pChem.exe*" pchem.exe .

The message "**Please press any key to continue**" appears when program runs to completion.

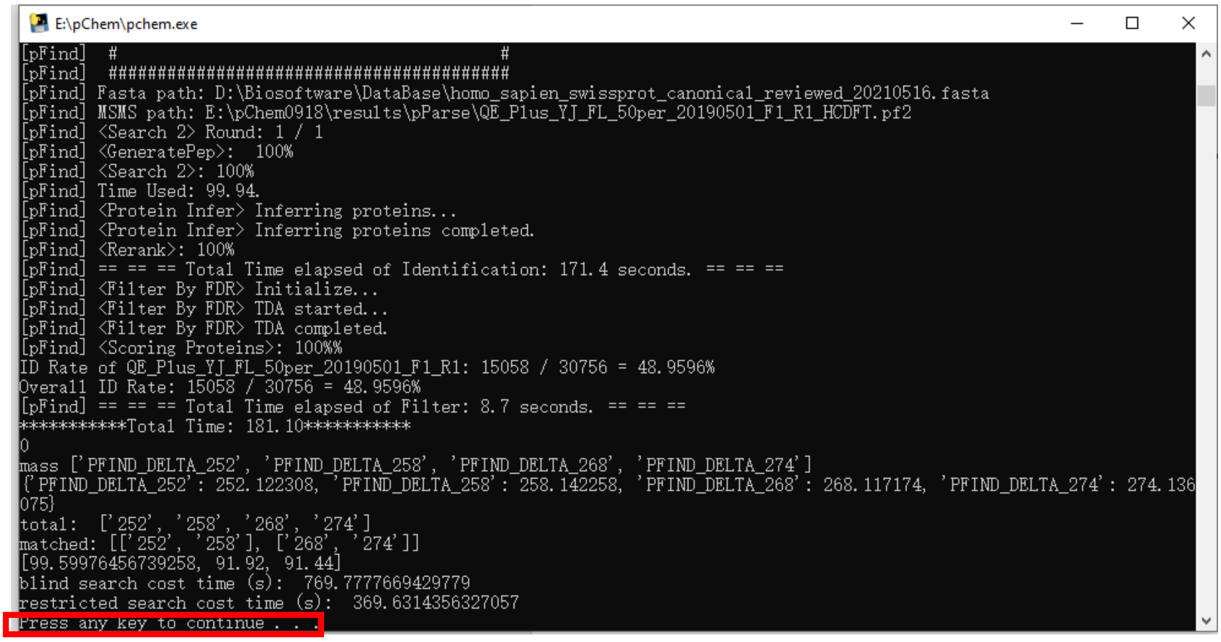

### 4. Output

4.1. Double click “*results*” file 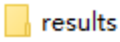

4.2. Double click “*reporting summary*”

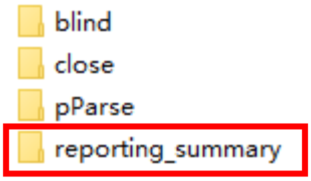

4.3. There are three major output documents.

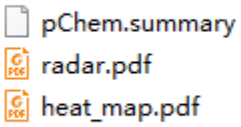

**Note:** Users are recommended to copy these output documents and paste into another file. Otherwise, they can be covered by those generated from the next search event.

#### ① pChem.summary

pChem.summary is a tab-delimited text file contains the details of each PDM candidate.

<table>
<tr><th>Rank</th><th>Mass Shift</th><th>Isotopic Label</th><th>Peptide Total</th><th>PSM Total</th><th>Peptide L|H</th><th>PSM L|H</th><th>Top1 Site</th><th>Top1 Probability</th><th>Accurate Mass</th><th>Site (Location /Proba</th><th>DFLs</th></tr>
<tr><td>1</td><td>PFIND_DELTA_252</td><td>Yes</td><td>5928</td><td>7350</td><td>3306|2622</td><td>4111|3239</td><td>C</td><td>0.94</td><td>252.121565|258.141564(22 ppm)</td><td>N-SIDE(0.111); A(0.0</td><td></td></tr>
<tr><td>2</td><td>PFIND_DELTA_268</td><td>Yes</td><td>478</td><td>644</td><td>250|228</td><td>352|292</td><td>C</td><td>0.412</td><td>268.117151|274.137027(42 ppm)</td><td>M(0.261); N-SIDE(0.2</td><td>302.109799, 320.119426,</td></tr>
</table>

Mass Shift: probe-derived modifications (Da)

Peptide Total: The number of modified peptides identified by blind search

PSM Total: The number of PSMs corresponding to modified peptides identified by blind search

Peptide L|H: The number of light and heavy peptides bearing the corresponding PDM, respectively

PSM L|H: The number of PSMs assigning to light and heavy bearing the corresponding PDM, respectively

Top1 site: The amino acid most likely to be labeled by the probe

Top1 Probability: $p_{site_i} = \frac{n_{site_i}}{n_{total}}$, In this formula, $p_{site_i}$ denotes the localization probability of PDM occurring $site_i$, $n_{site_i}$ is the number of

PSMs related to each PDM that occur on specific $site_i$, and $n_{total}$ denotes the total number of PSMs related to the same PDM.

Site: Other amino acid sites that may also be labeled by probes and their corresponding localization probability values

DFLs: Diagnostic fragment losses

② **Heat_map.pdf**

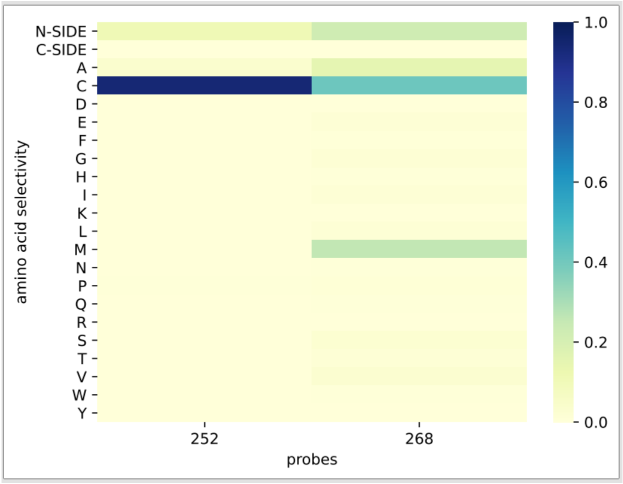

Horizontal coordinate: The Δmass of each PDM

Longitudinal coordinate: The types of amino acids

Color gradient: The localization probability that the modification occurs at each potential site.

③ **Radar.pdf**

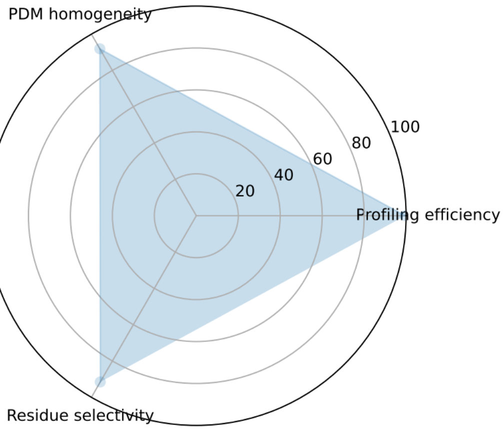

Radar.pdf contains a radar plot whose radial axes correspond to the three features/scores calculated based on the values extracted from **pChem.summary** as follows.

| Rank | Mass Shift | Isotopic Label | Peptide Total | PSM Total | Peptide L\|H | PSM L\|H | Top1 Site | Top1 Probability | Accurate Mass | Site (Location /Probability)-Others |
|---|---|---|---|---|---|---|---|---|---|---|
| 1 | PFIND_DELTA_Δ1 | | | A | | | Cys | a | | |
| 2 | PFIND_DELTA_Δ2 | | | B | | | lys | b | | Cys/e |
| … | … | | | … | | | … | … | | |
| N | PFIND_DELTA_ΔN | | | X | | | … | x | | |

PDM homogeneity: A/( A+B+…+X)

Residue selectivity: (a·A+b·e)/( A+B+…+X)

Profiling efficiency: #PSMs for the PDMs shown in pChem.summary/ #All PSMs

**Note:**
PDM homogeneity and Residue selectivity are calculated based on the blind search results. Profiling efficiency is calculated according to restricted search (see "pChem.close.summary" file that located in "X:\pChem\results\close\pChem.close.summary")

#### 5. Supporting protocol 1

How to download protein *.fasta files for database search?

5.1. Open https://www.uniprot.org/, enter the Latin name of the species (e.g., *home sapiens*), then click search

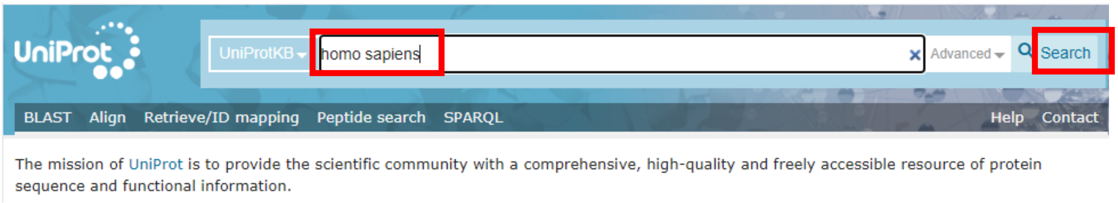

5.2. Click "*Reviewed*" (Swiss-Prot)

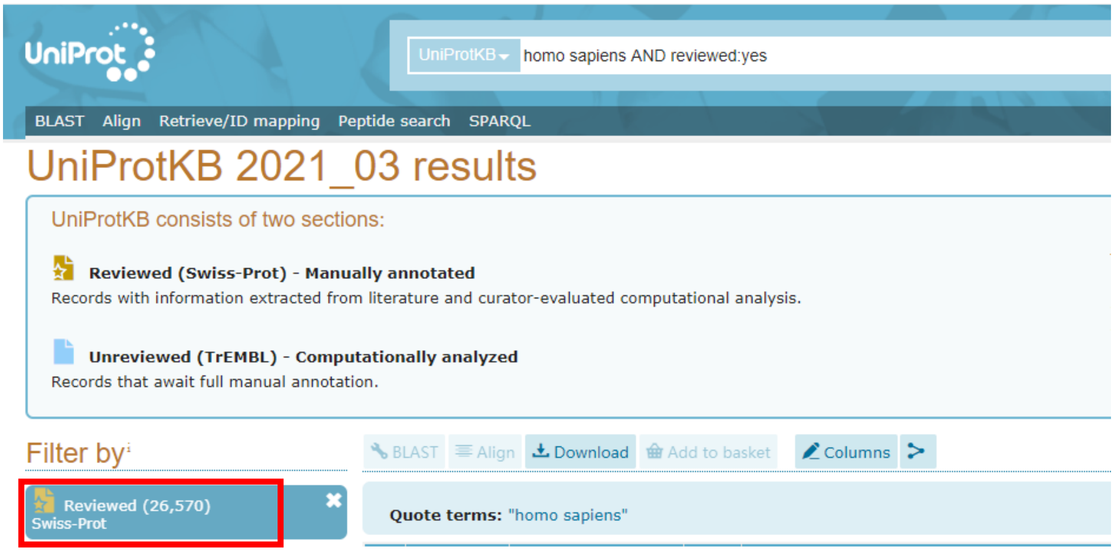

5.3. Select "*Uncompressed*" , then Click "*Download*" and "*Go*"

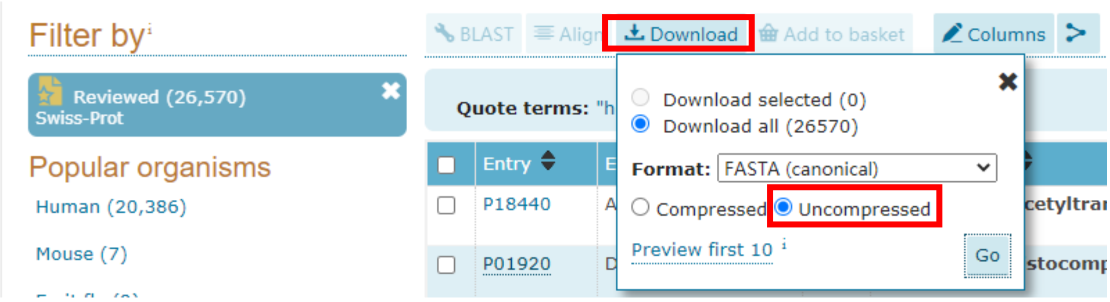

5.4. Get the *.fasta file

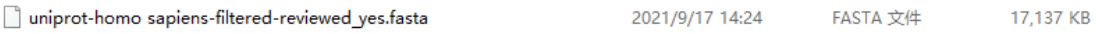

#### 6. Supporting protocol 2

How to convert non-Thermo MS data into mzML format files for pChem search?

6.1. Download MSconvertGUI that is embedded in the ProteoWizard platform from: [https://proteowizard.sourceforge.io/download.html](https://proteowizard.sourceforge.io/download.html)

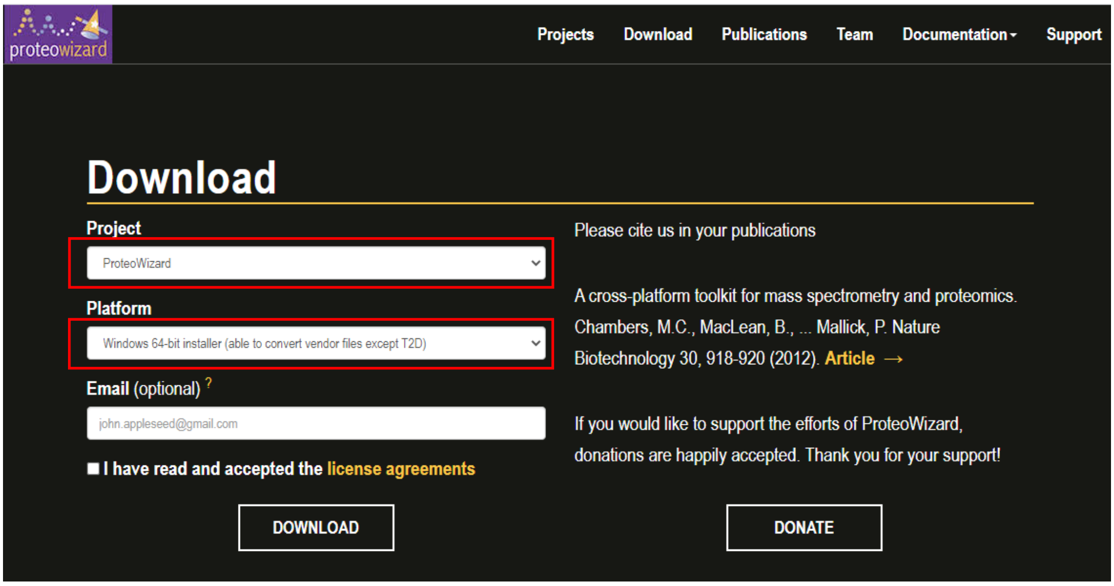

6.2. Install ProteoWizard according to the following instruction

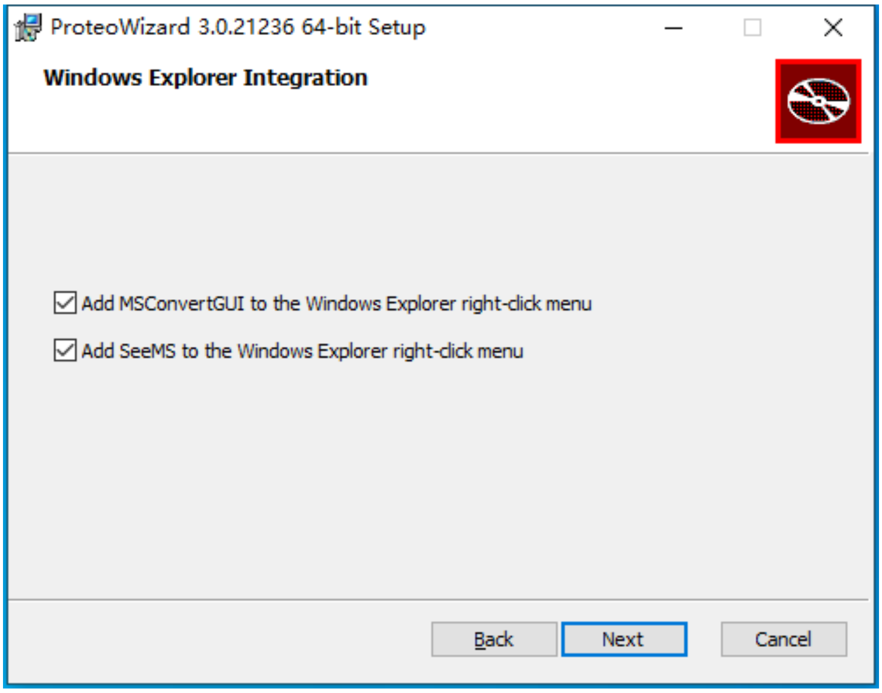

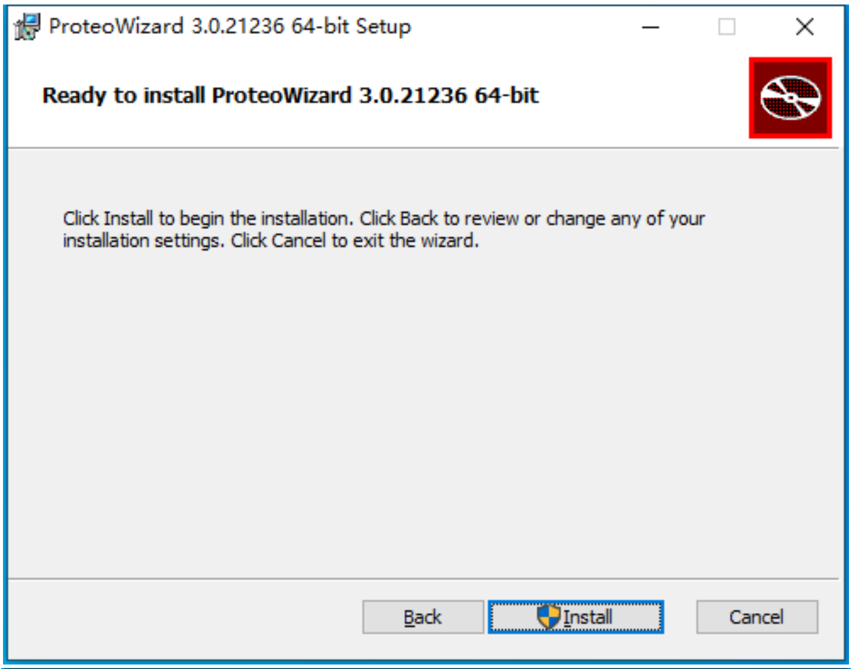
ProteoWizard 3.0.21236 64-bit Setup
Ready to install ProteoWizard 3.0.21236 64-bit
Click Install to begin the installation. Click Back to review or change any of your installation settings. Click Cancel to exit the wizard.
Back
Install
Cancel

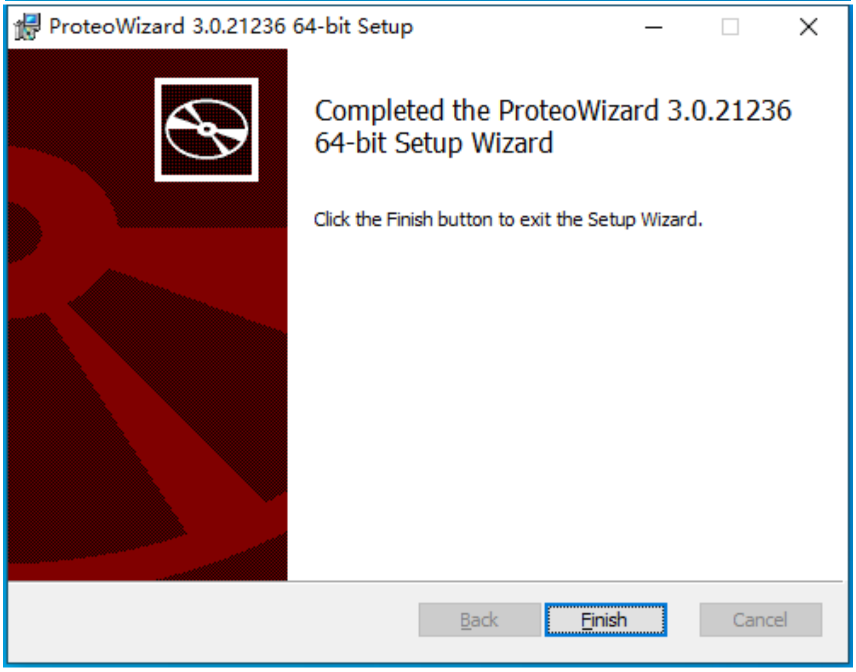
ProteoWizard 3.0.21236 64-bit Setup
Completed the ProteoWizard 3.0.21236 64-bit Setup Wizard
Click the Finish button to exit the Setup Wizard.
Back
Finish
Cancel

#### 6.3. Open MSconvertGUI

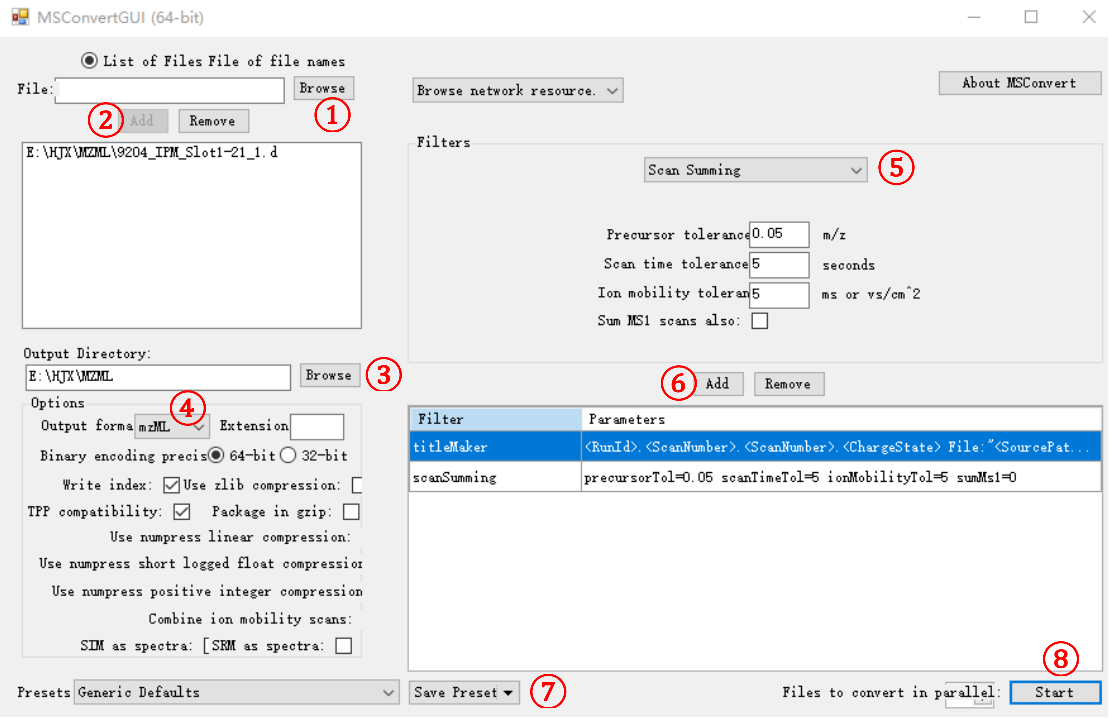

①-② Browse and add MS data (e.g., *.d, *.WIFF files)

③ Define output route

④ Choose *.mzML as the output data format

⑤-⑥ Define parameters for Scan Summing

⑦-⑧ Save and run
