## Supplementary Information for "pChem: a modification-centric assessment tool for the performance of chemoproteomic probes"

### SUPPLEMENTARY NOTES

#### SUPPLEMENTARY NOTE 1：Processing products of IPM-derived modifications

The mass shift of Δ268.12 on cysteine or methionine and the targeted PDM by IPM differ by 16.0 Da, which is corresponding to one oxygen atom (**Supplementary Fig. 2a**). Since, in a chemical aspect, it is unlikely for IPM to react with methionine, Δ268.12 may be explained by the co-occurrence of methionine oxidation and the targeted PDM on cysteine, which cannot be distinguished by the MS/MS information due to the lack of modification-specific fragment ions. In this regard, we found that Δ268.12 is often assigned to methionine when it is adjacent to a cysteine, and that such PSMs prone to yield a classic oxidation[M]-derived neutral loss of 64.0 Da (**Supplementary Fig. 2b**). This finding further ruled out the possibility of Δ268.12 on methionine. Moreover, targeted search revealed that 91.5% PSMs bearing Δ268.12 were re-annotated to pinpoint the targeted PDM on cysteine. Note, similar miss-matching was also observed for the cysteine-targeting probes. On the other hand, Δ268.12 on cysteine can be a sulfoxide product derived from the targeted PDM of Δ252.12, which is further supported by the DFLs (**Supplementary Table 2**). Notably, we found that Δ268.12 also occurred on a non-Cys peptide of MIF protein (**Supplementary Fig. 2c**). This initially puzzling outcome was explained when we noted that this peptide contains a catalytic N-term proline with a high nucleophilicity (i.e., with an unusually low pKa of 5.6[1]) and an adjacent methionine.

### SUPPLEMENTARY NOTE 2：Expanding the aHNE/aONE-derived adductomes

Using alkyne surrogates of HNE and ONE (i.e., aHNE and aONE, respectively), we previously profiled dozens to hundreds of adducted sites by these two lipid electrophiles[2, 3]. We also designed a strategy to determine the turnover of aHNE/aONE-derived adducts in intact cells (**Supplementary Fig. 14a**). Specifically, cells were first labeled with aHNE or aONE. After 2 h of incubation, cells were either harvested immediately or placed in the probe-free medium for another 1 and 4 h recovery period, respectively, and processed into peptide samples for LC-MS/MS analyses (See **Methods**). By re-searching the resulting data sets with two additional PDMs, here we identified 6.2% and 23.4% more adducted sites for aHNE and aONE, respectively (**Supplementary Fig. 14b-c**). Quantitative analyses further confirmed the fast turnover rates of most adduction events, while revealed a high stability of GSH-aONE-lysine conjugate at 1 and 4 h of recovery (**Supplementary Fig. 14d-e**).

#### References


1. Stamps, S.L., Fitzgerald, M.C. & Whitman, C.P. Characterization of the role of the amino-terminal proline in the enzymatic activity catalyzed by macrophage migration inhibitory factor. *Biochemistry* **37**, 10195-10202 (1998).
2. Yang, J., Tallman, K.A., Porter, N.A. & Liebler, D.C. Quantitative chemoproteomics for site-specific analysis of protein alkylation by 4-hydroxy-2-nonenal in cells. *Anal Chem* **87**, 2535-2541 (2015).
3. Sun, R. et al. Chemoproteomics Reveals Chemical Diversity and Dynamics of 4-Oxo-2-nonenal Modifications in Cells. *Mol Cell Proteomics* **16**, 1789-1800 (2017).

### SUPPLEMENTARY FIGURES

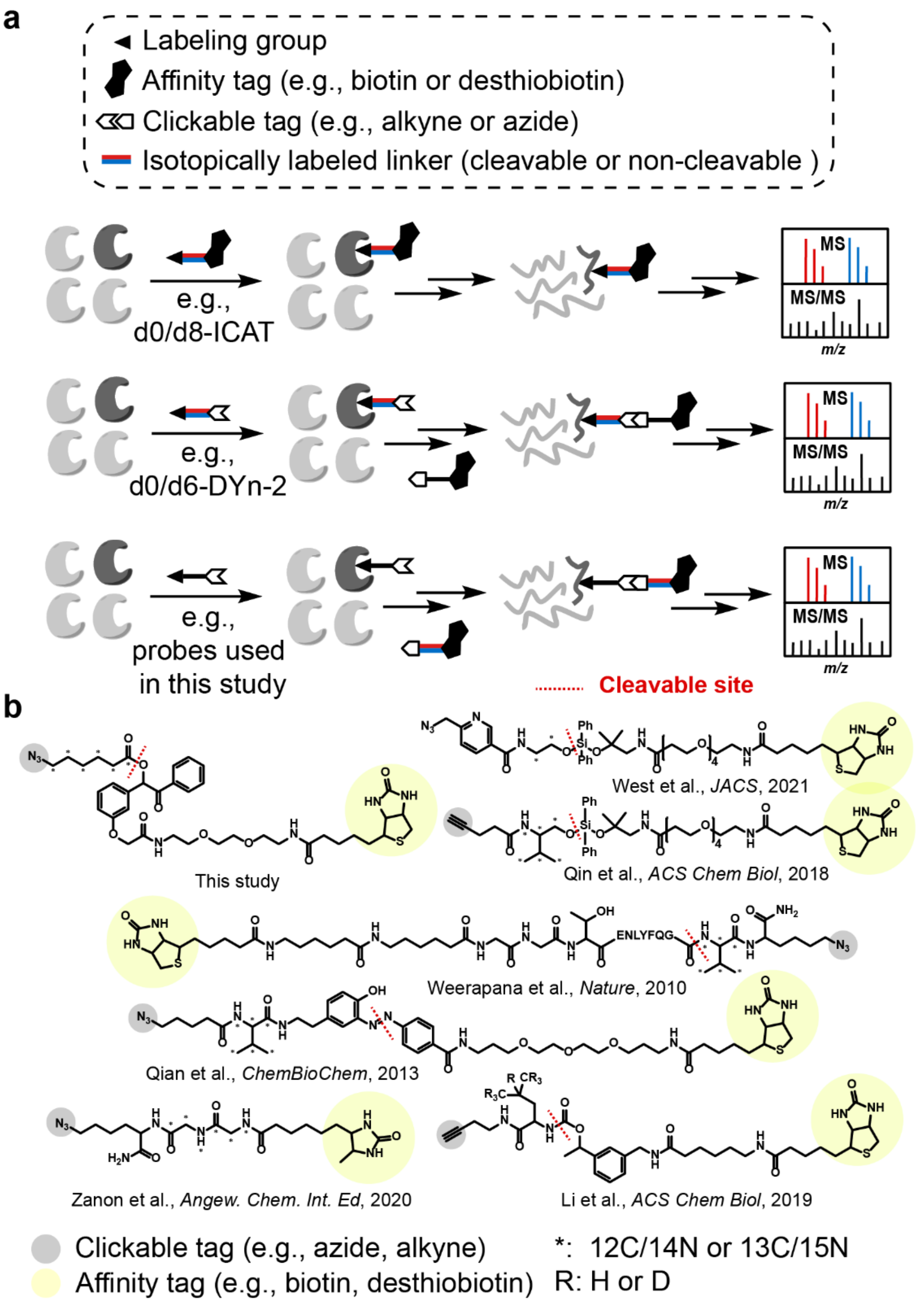


**Supplementary Fig. 1. General strategies for isotope-coding of probe-derived modifications.** **a**, Schematic of the workflows for quantitative chemoproteomics. *Upper*, the 'enrichable' probes with an affinity tag are isotopically coded, such as the well-known d0/d8 ICAT reagents. *Middle*, the 'clickable' probe itself is isotopically coded, such as d0/d6 DYn-2; *Bottom*, the 'clickable' probe is first used for proteomic labeling and its PDMs can then be conjugated with isotopically labeled linkers. **b**, Isotope-coding reagents for 'clickable' probes (i.e., isotopically labeled affinity linkers) used in this and many other studies.

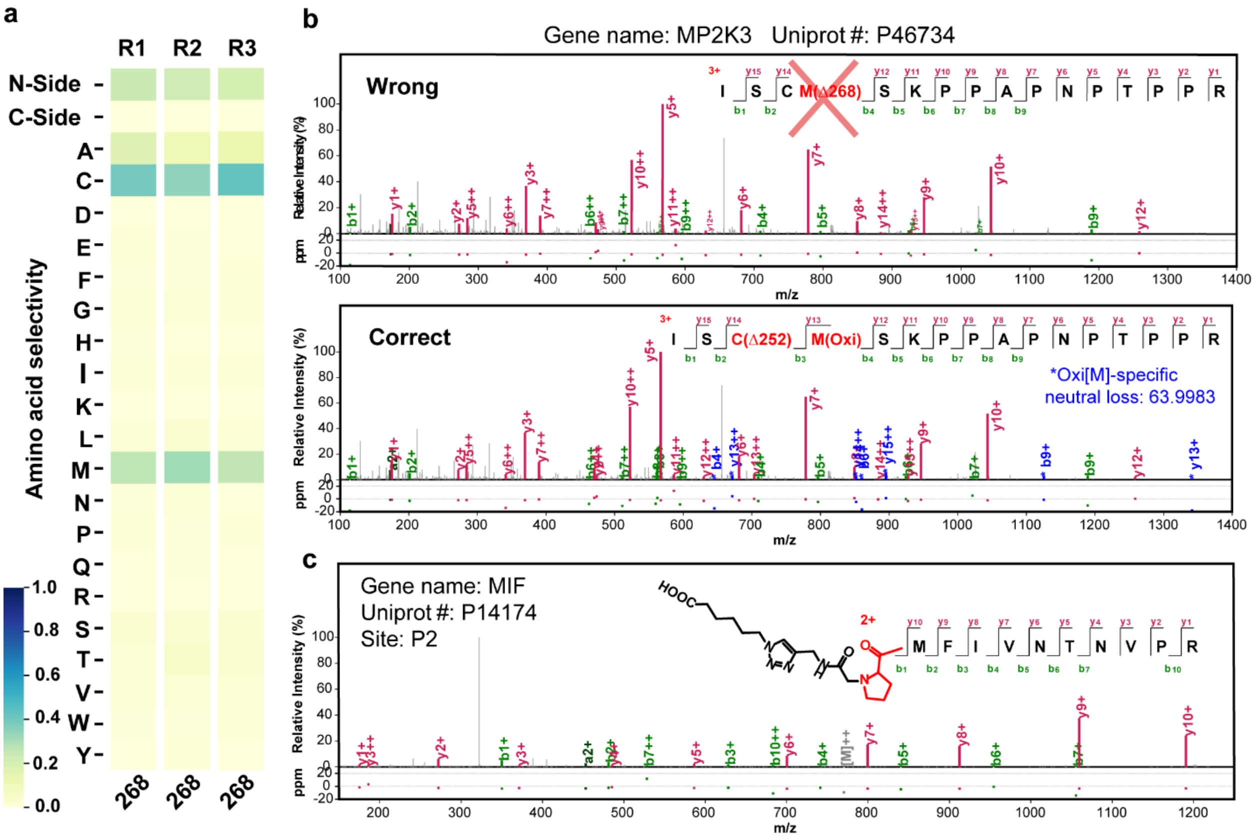


**Supplementary Fig. 2. Identification of an unexpected but minor modification that is derived from IPM. a**, Heatmaps showing the amino acid localization distribution of the IPM-derived modification of Δ268.12. **b**, Representative MS/MS spectrum wrongly assigning the targeted PDM (Δ252.12) to oxidized methionine due to the presence of adjacent cysteine. **c**, Representative MS/MS spectrum of a peptide bearing the PDM Δ268.12 assigning to the protein N-term proline of MIF.

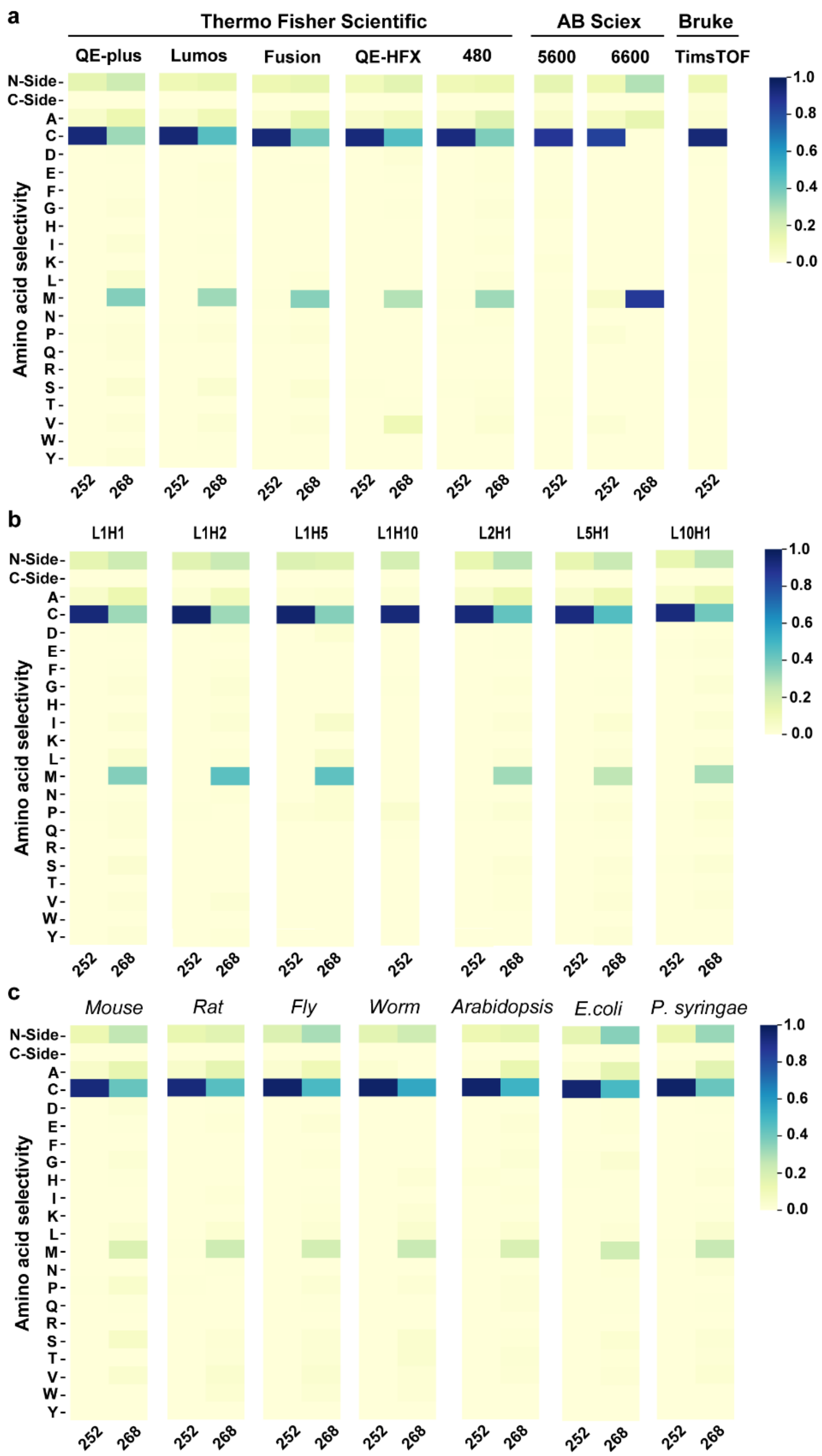


**Supplementary Fig. 3. Robustness of pChem.** Representative heatmaps generated by pChem showing the amino acid localization distribution of the IPM-derived modifications. **a**, IPM-based QTRP data for pChem search were produced on eight different LC-MS/MS instruments from three vendors. **b**, IPM-based QTRP data for pChem search were produced previously from light and heavy IPM-tagged samples mixed in different ratios (L/H = 1:10, 1:5, 1:2, 1:1, 2:1, 5:1 and 10:1). **c**, Data for pChem search were produced from the IPM-based QTRP applications in various species.

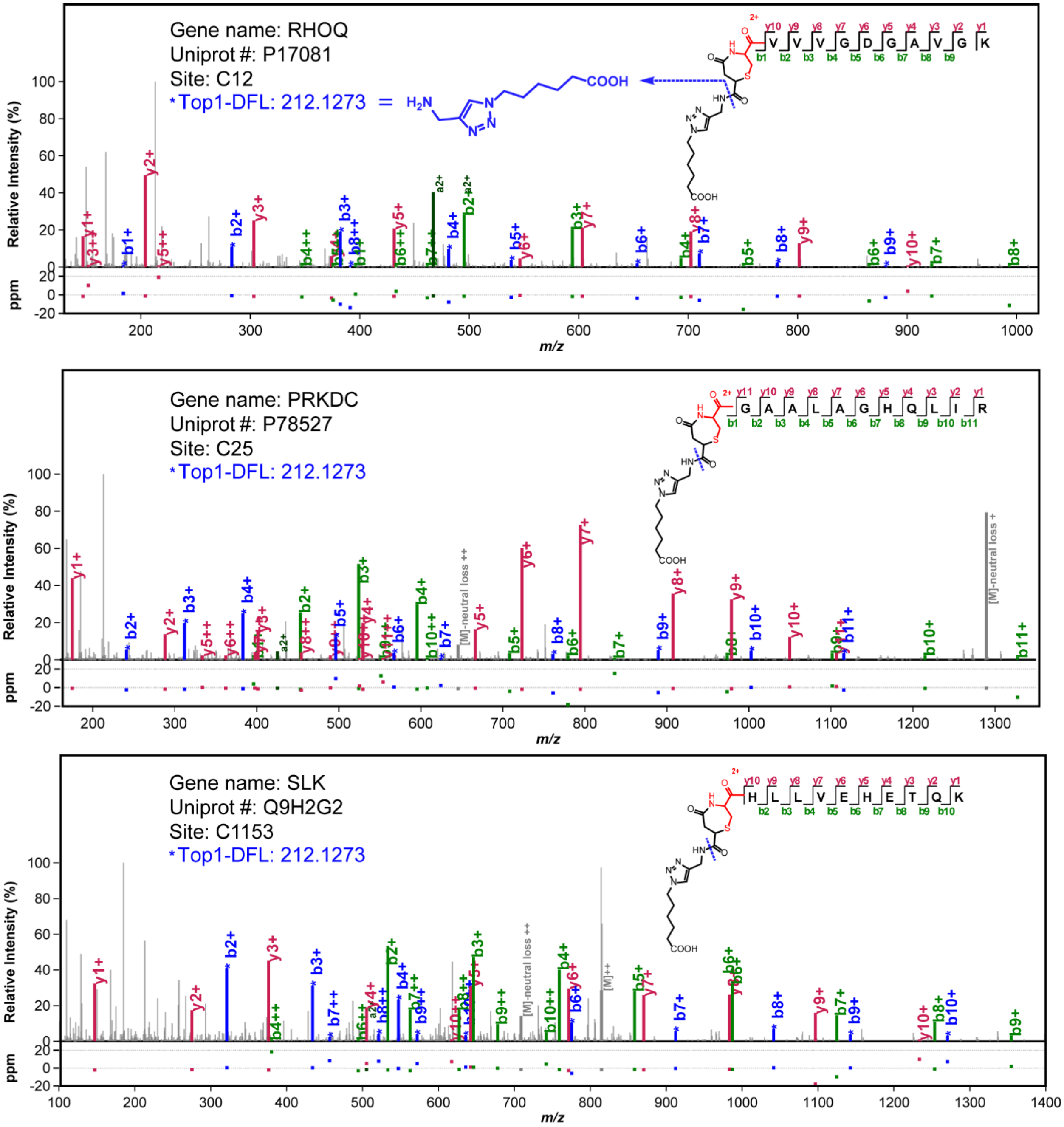


**Supplementary Fig. 4. Characterization of a previously unknown peptide N-term cysteine modification (Δ292.12) derived from NPM.** Representative MS/MS spectra of the peptides bearing this PDM on their N-term cysteines. Sequence fragment ions with the top1 diagnostic fragment loss (DFL) are annotated in blue color. This DFL is generated from the cleavage of linear amide bond on the PDM (blue dash line).

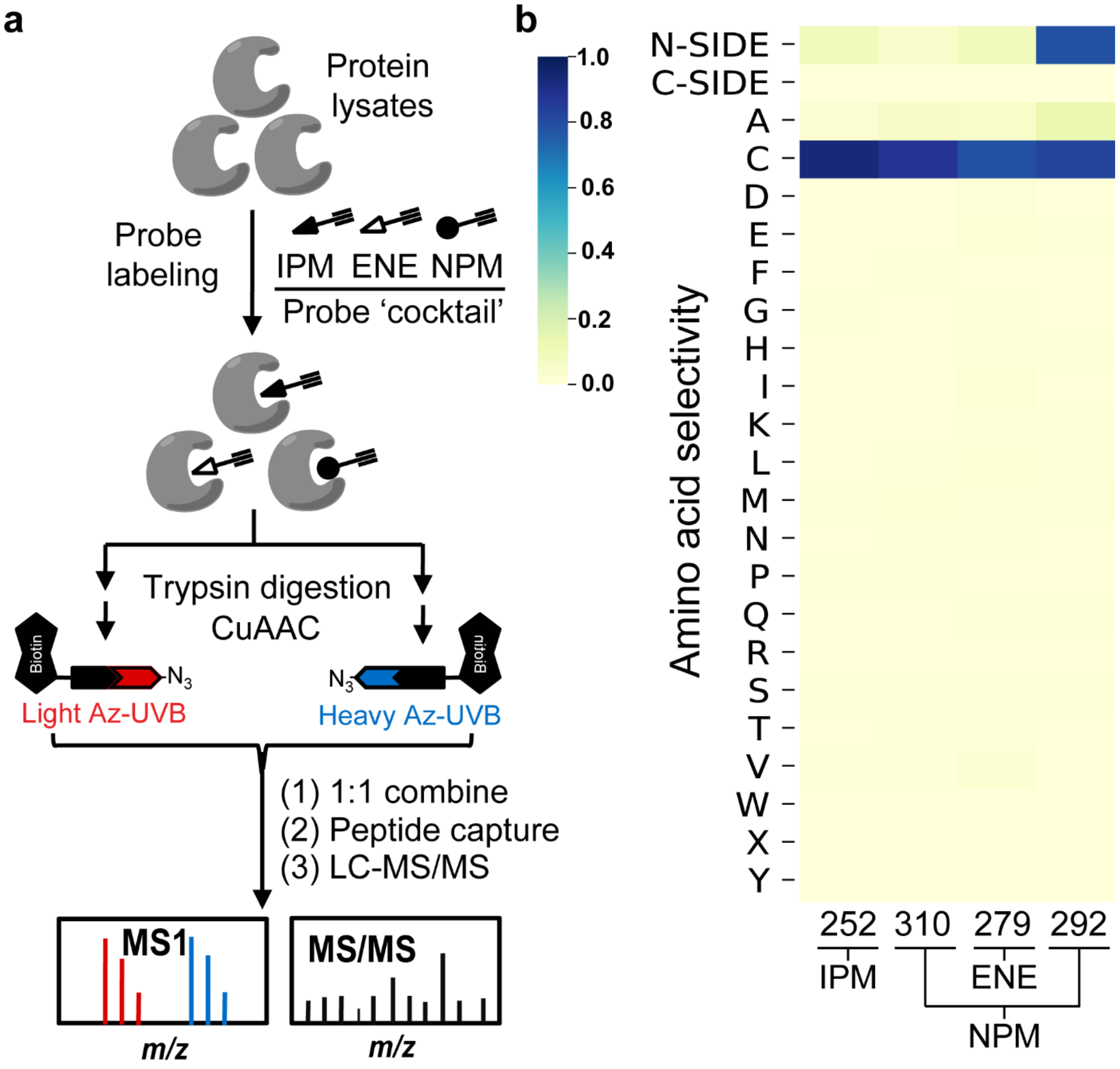


**Supplementary Fig. 5. Benchmarking pChem with a "cocktail" dataset. a**, Schematic of the experiment design for generating a QTRP dataset using a 'cocktail' of three thiol-reactive probes, including IPM, ENE, and NPM. **b**, Representative heatmaps showing the amino acid localization distribution of the pChem-defined PDMs.

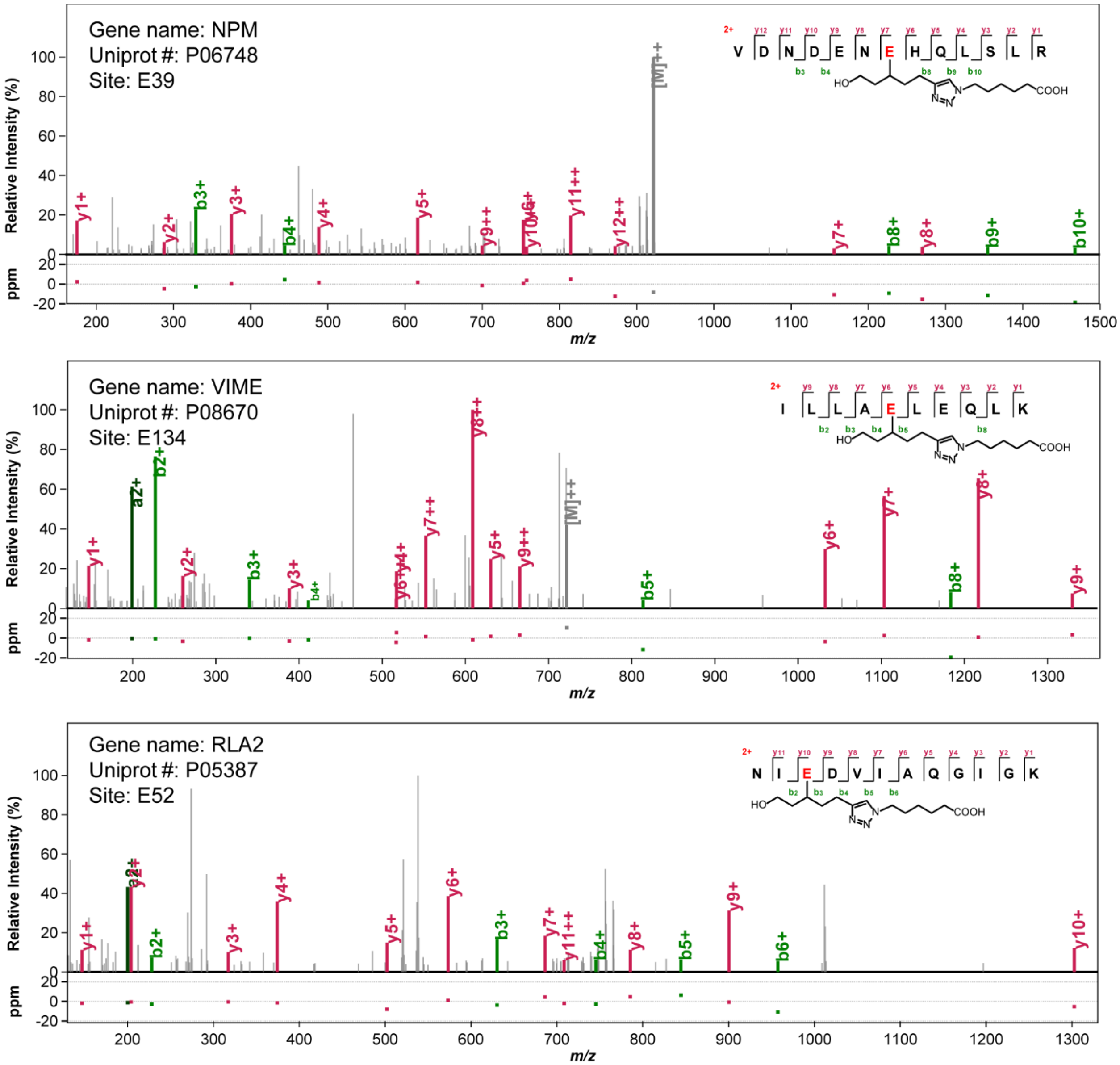


**Supplementary Fig. 6. Characterization of the glutamic acid-preference of a diazirine-based probe.** Representative MS/MS spectra of the peptides bearing the PDM of Δ267.16 on glutamic acid.

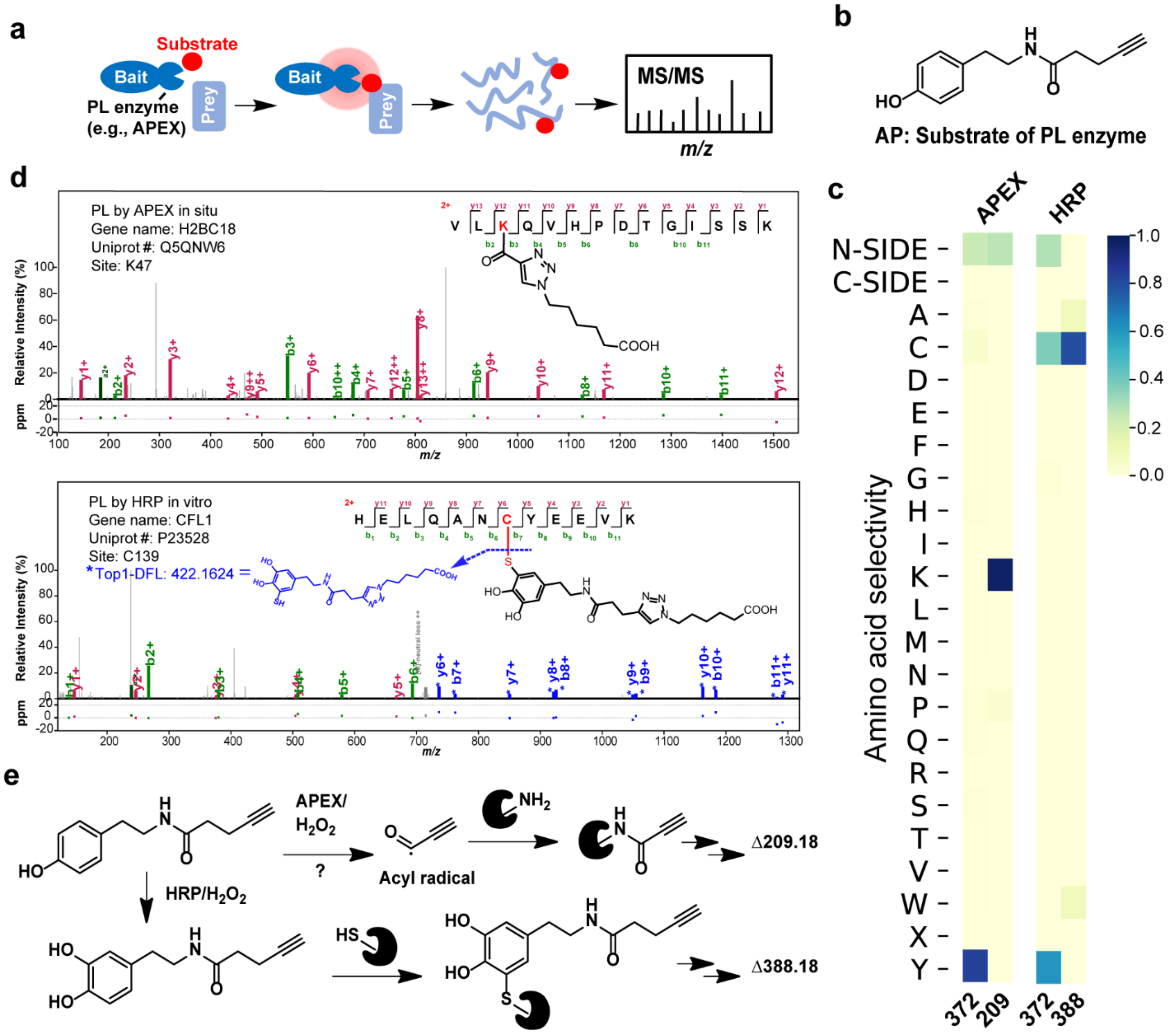


**Supplementary Fig. 7. Application of pChem for AP, a substrate probe of proximity labeling enzyme. a**, Schematic of peroxidase-based proximity labeling. A peroxidase itself or its fusion with a bait protein catalyzes its substrate (e.g., alkyne phenol, AP) to covalently modify proximal prey proteins. Probe labeled peptides can be enriched for LC-MS/MS analysis, followed by site-level identifications of prey proteins. **b**, Chemical structure of AP. **c**, Representative heatmap showing the amino acid localization distribution of the AP-derived modifications in two different experimental settings. **d-e**, Characterization of selected PDMs from AP. *Upper*, a representative MS/MS spectrum of peptide bearing the PDM of Δ209.18 on lysine. *Lower*, a representative MS/MS spectrum of peptide bearing the PDM of Δ388.18 on cysteine. Sequence fragment ions with the top1 diagnostic fragment loss (DFL) are annotated in blue color. This DFL is generated from the cleavage of C-S bond on the PDM (blue dash line). **e**, Plausible mechanisms for the formation of two AP-based PDMs on lysine and cysteine, respectively.

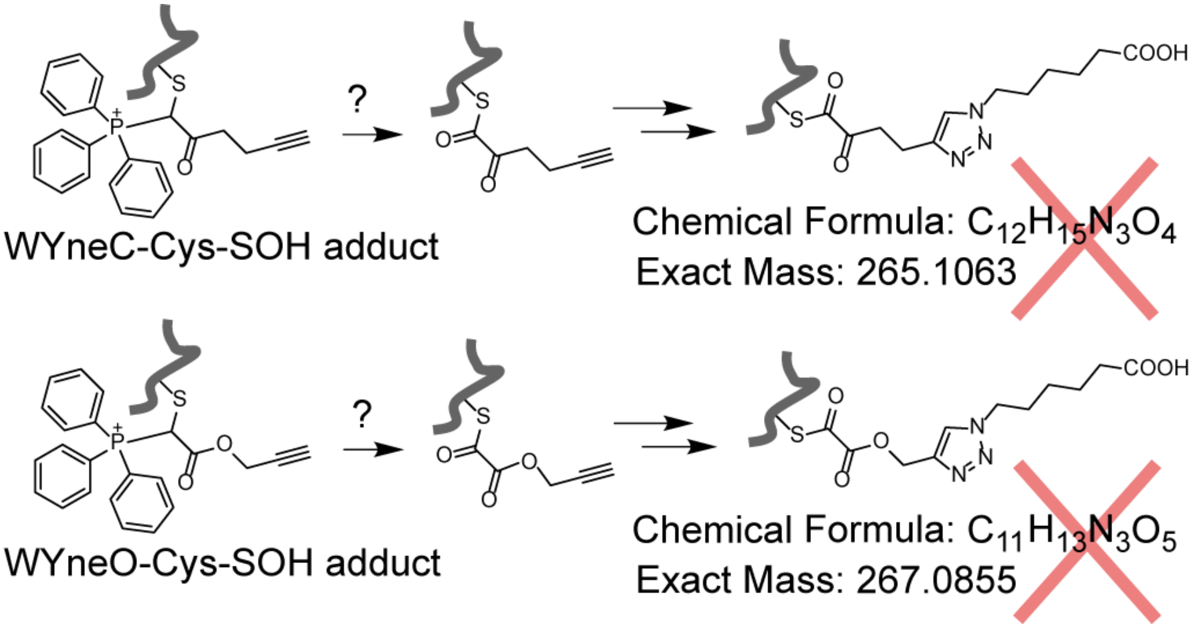


**Supplementary Fig. 8. The previously reported mechanism for the formation of TPP-cleaved peptide adducts derived from WYneC/O**

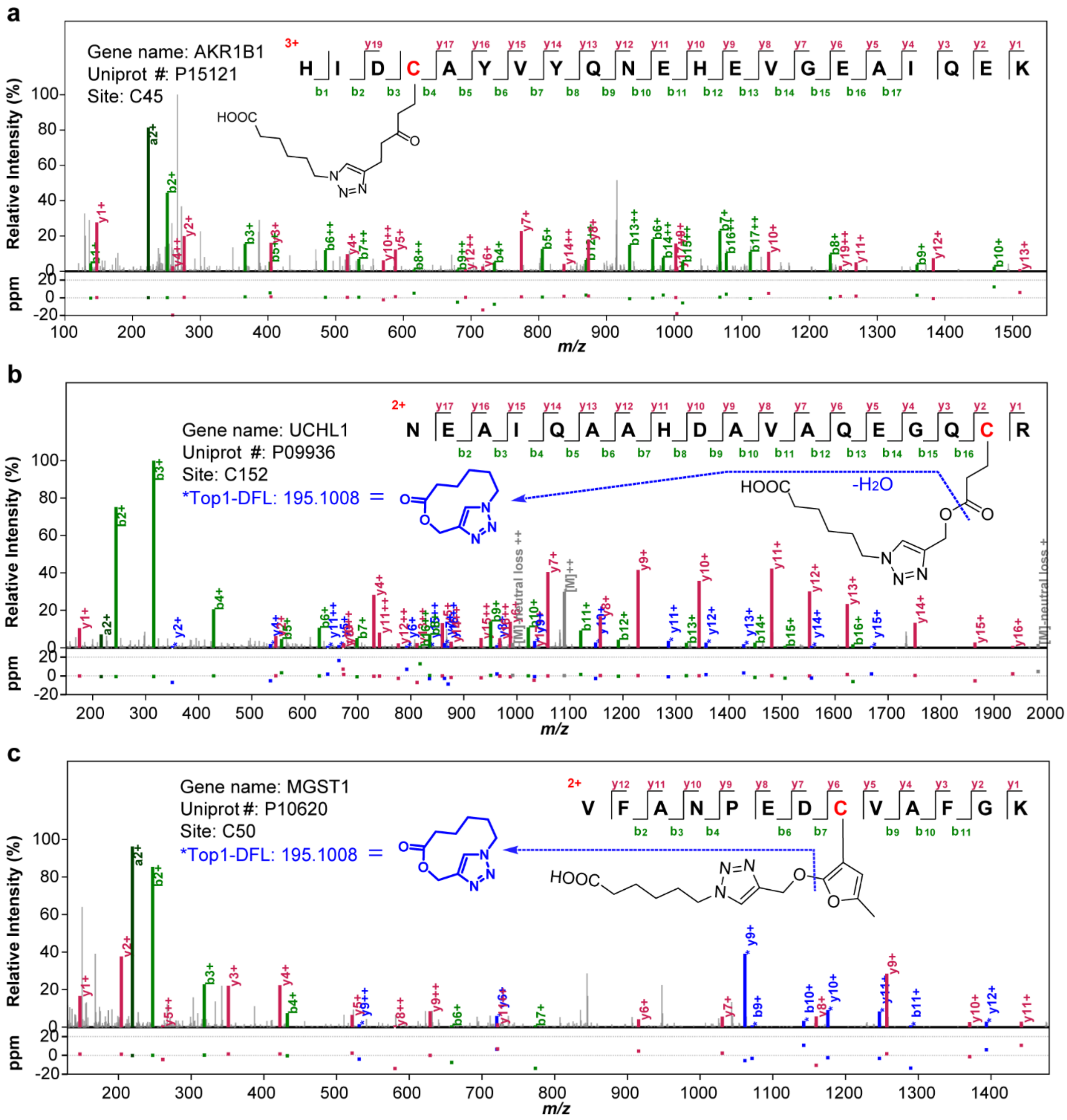


**Supplementary Fig. 9. Characterization of triphenylphosphonium (TPP)-loss modifications derived from WYneC/O. a**, Representative MS/MS spectrum of a peptide bearing the PDM of Δ265.14 from WYneC. **b**, Representative MS/MS spectrum of a peptide bearing the PDM of Δ267.12 from WYneO. **c**, Representative MS/MS spectrum of a peptide bearing the PDM of Δ291.12 from WYneO. For **b-c**, sequence fragment ions with the top1 diagnostic fragment loss (DFL) are annotated in blue color. These DFLs are generated from the cleavage of ester bond on the corresponding PDMs (blue dash line).

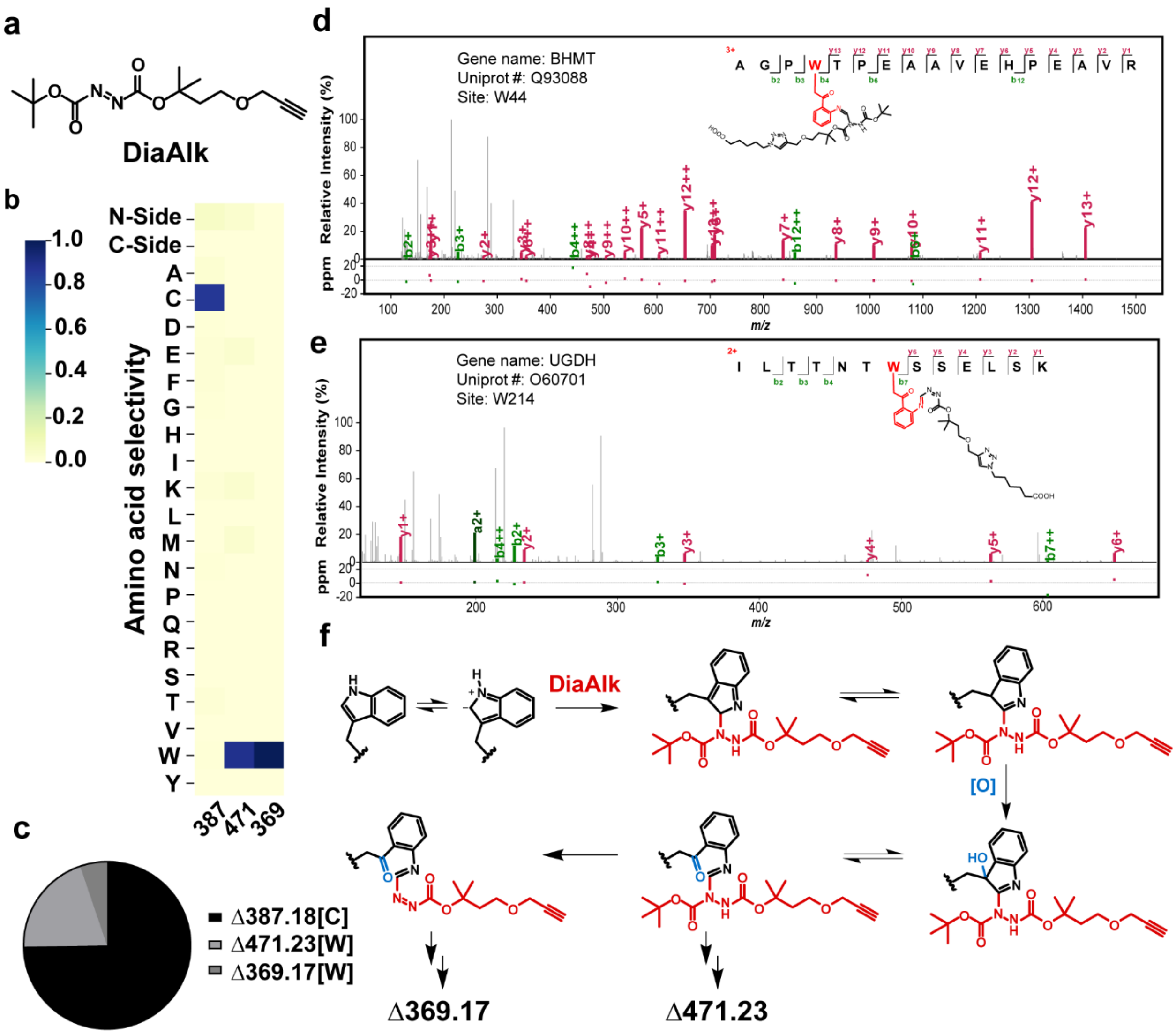


**Supplementary Fig. 10. Application of pChem for DiaAlk, a $SO_2H$ probe. a**, Chemical structure of DiaAlk. **b**, Representative heatmaps showing the amino acid localization distribution of the pChem-defined PDMs for DiaAlk. **c**, Pie charts showing the abundance distribution (i.e., number of PSMs) of PDMs from DiaAlk. **d**, Representative MS/MS spectrum of a peptide bearing the PDM of Δ471.23 on tryptophan. **e**, Representative MS/MS spectrum of a peptide bearing the PDM of Δ369.17 on tryptophan. **f**, Plausible mechanism for the formation of two tryptophan-targeting PDMs (i.e., Δ471.23 and Δ369.17) both derived from DiaAlk.

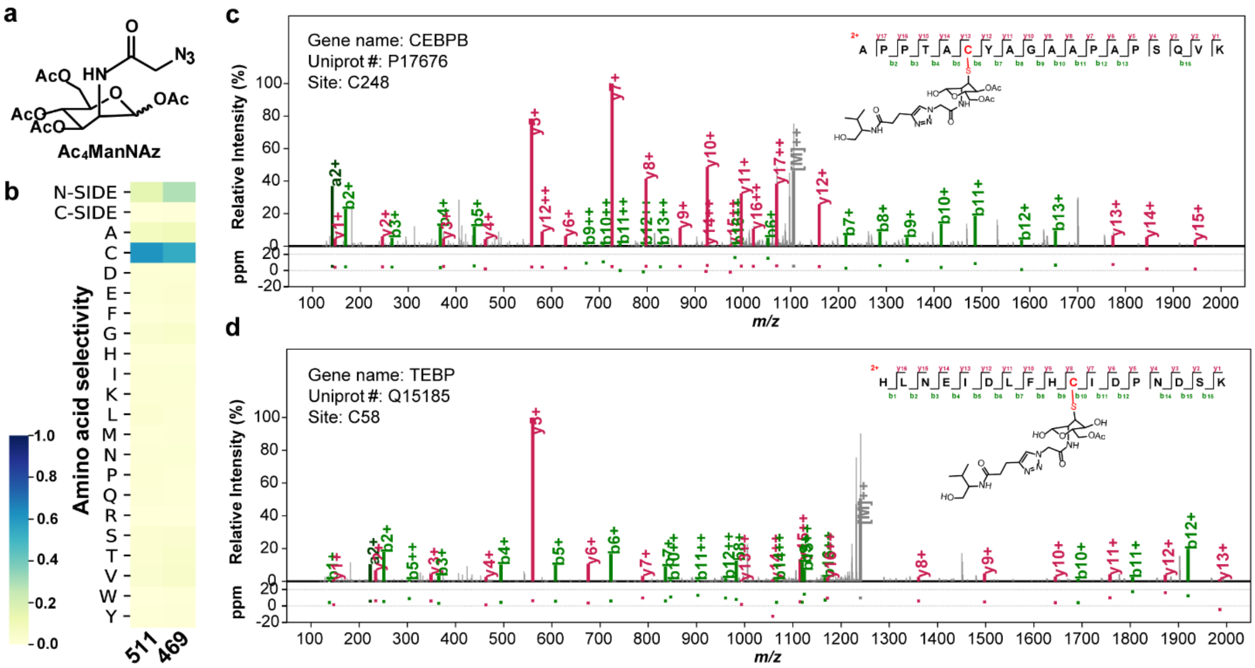


**Supplementary Fig. 11. Application of pChem for $Ac_4ManNAz$, a metabolic labeling probe for glycoproteomics. a**, Chemical structure of $Ac_4ManNAz$. **b**, Representative heatmaps showing the amino acid localization distribution of the pChem-defined PDMs for $Ac_4ManNAz$. **c**, Representative MS/MS spectrum of a peptide bearing the PDM of Δ511.23 on cysteine. **d**, Representative MS/MS spectrum of a peptide bearing the PDM of Δ469.22 on cysteine.

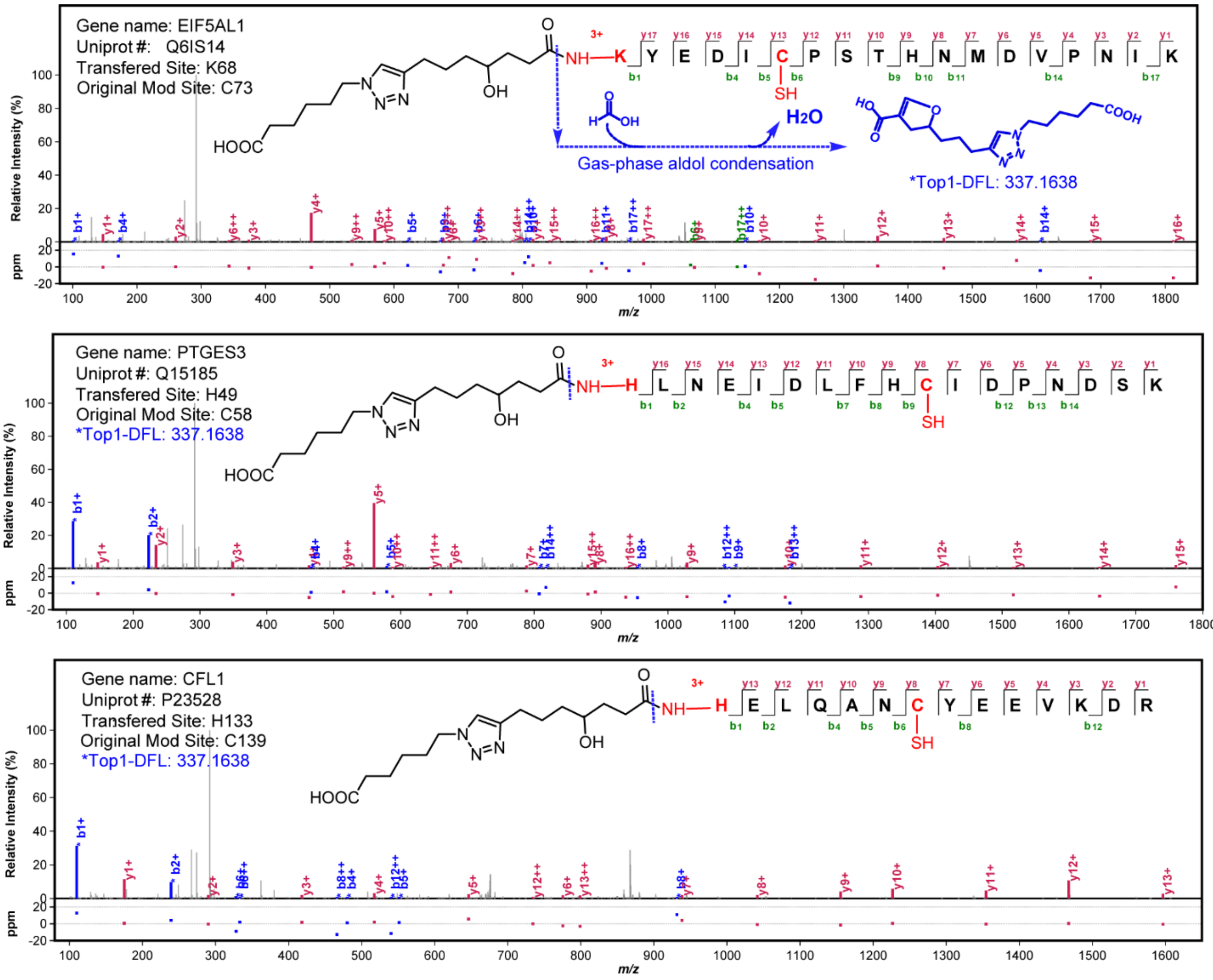


**Supplementary Fig. 12. Characterization of a peptide N-term modification (∆309.17) derived from aHNE.** Representative MS/MS spectra of peptides bearing this PDM on sequence N-term. Sequence fragment ions with the top1 diagnostic fragment loss (DFL) are annotated in blue color. This DFL is generated from the cleavage of amide bond on the PDM, immediately followed by the aldol condensation reaction with formic acid in gas-phase (blue dash line).

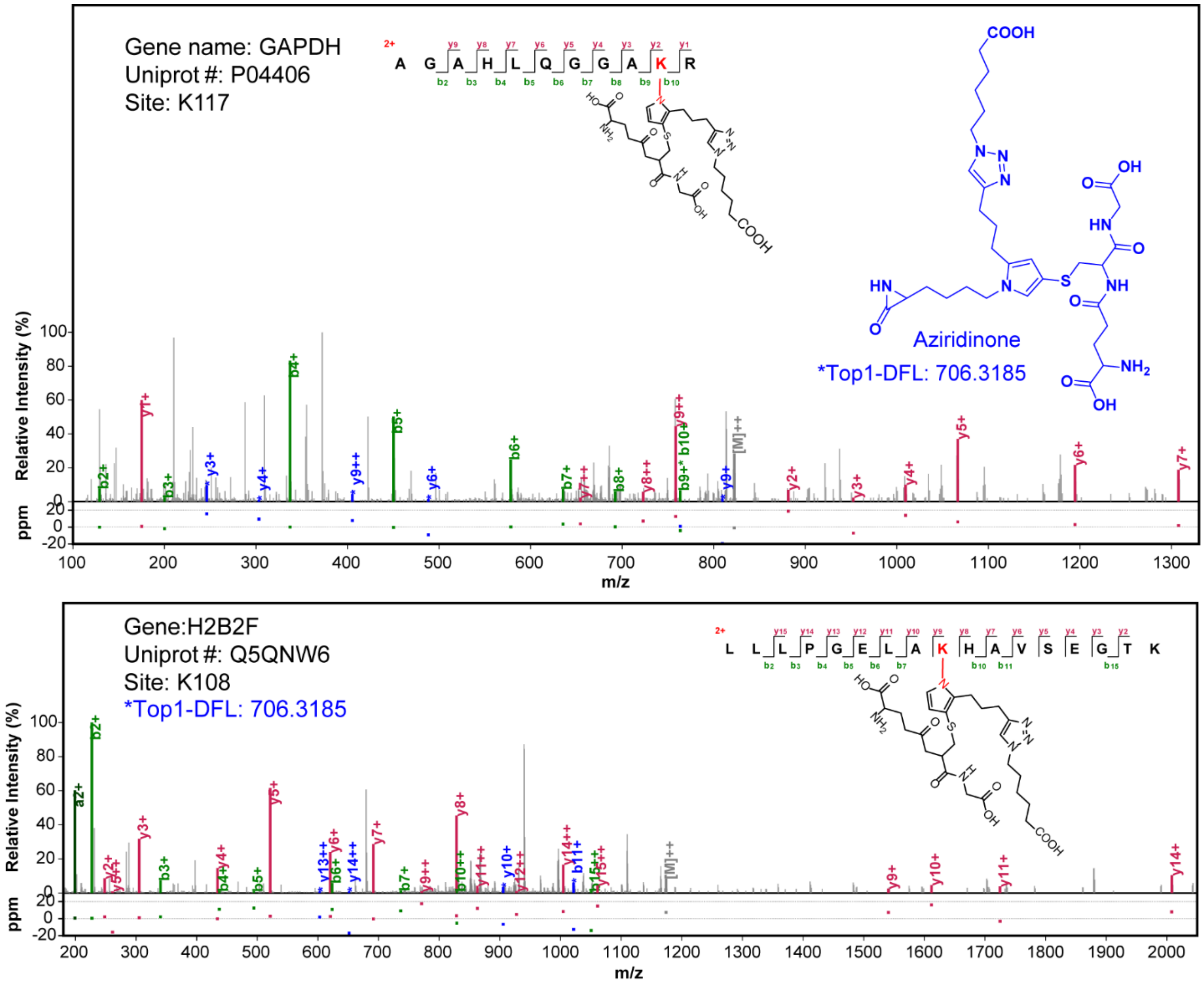


**Supplementary Fig. 13. Characterization of a lysine modification (∆578.22) derived from aONE.** Representative MS/MS spectra of peptides bearing this PDM on lysine. Sequence fragment ions with the top1 diagnostic fragment loss (DFL) are annotated in blue color. This DFL is generated from the cleavage of amide bond on the peptide sequence, resulting in the lysine-specific loss of aziridinone product (blue).

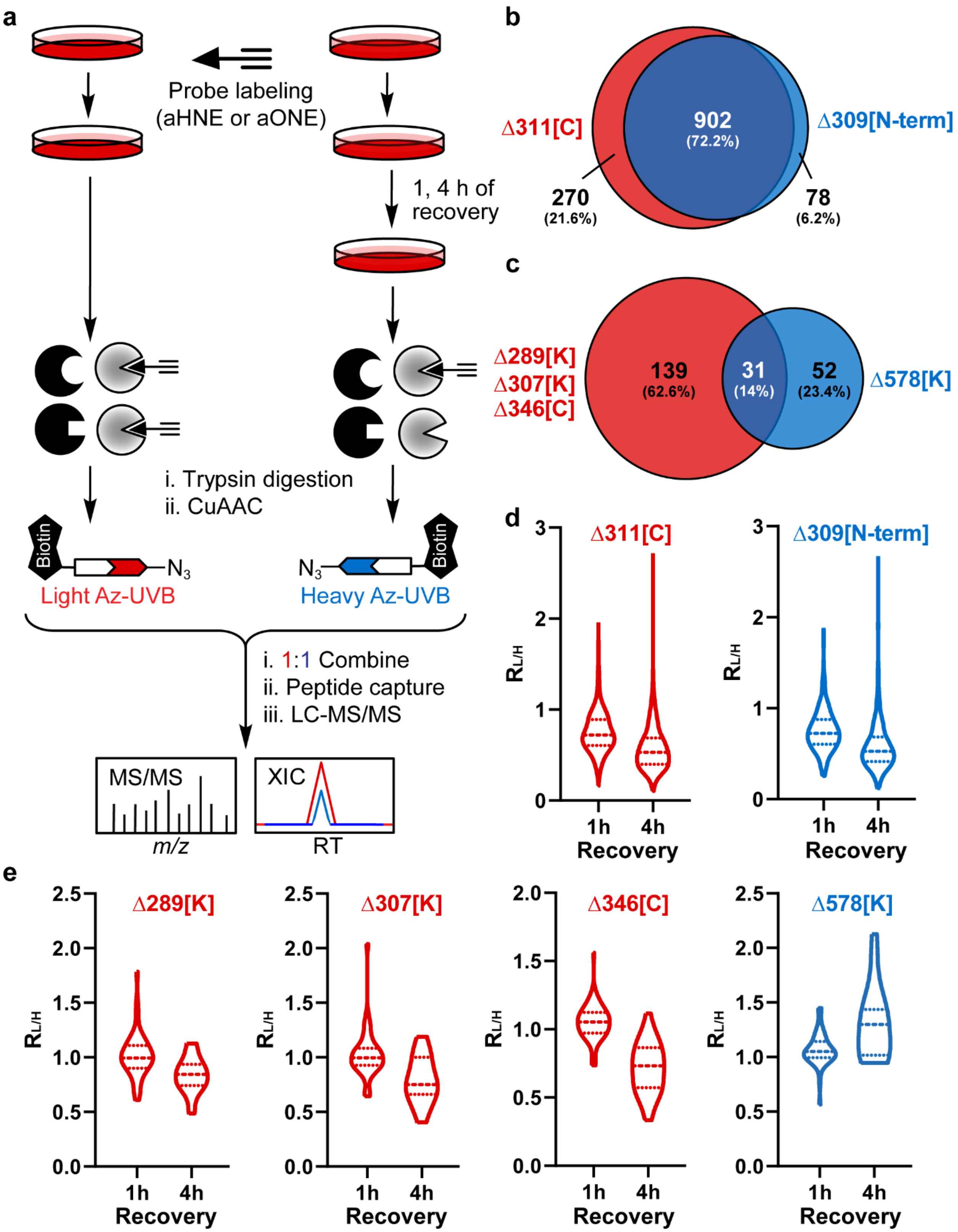


**Supplementary Fig. 14. Re-analyses of protein adduction by aHNE and aONE in RKO cells.** Raw data sets for such re-analyses were retrieved from Ref. [55] (Yang, et al., *Anal Chem*, 2015) and Ref. [24] (Sun, et al., *Mol Cell Proteomics*, 2017). **a**, Schematic of the workflow for quantitative chemoproteomic analyses of dynamic aHNE/aONE-derived protein adducts in RKO cells. Cells were first treated with either aHNE or aONE. After treatment, cells were either harvested immediately and used as controls or placed in probe-free medium for another 1 and 4 h recovery period. The probe-labeled proteomes were digested with trypsin and then biotinylated by click chemistry with the light (L, recovery) or heavy (H, control) labeled UV-cleavable azido biotin, followed by streptavidin enrichment, photorelease, and LC-MS/MS analysis. Identification and quantification were performed using the pFind studio (See **Methods** for more details). **b-c**, Venn diagrams revealing that pChem-based identification of previously unknown PDMs substantially expanded the target spectrum of aHNE (**b**) and aONE (**c**). Note, cysteines on those N-term ketoamide peptide adducts (Δ307.15) are assigned as aHNE-modified sites, since such a PDM is most likely generated through an intramolecular rearrangement from Cys to N-term. **d-e**, Dynamics of aHNE- and aONE-based PDMs in RKO cells. **d**, Violin plots of L/H ratios determined from two types of aHNE-derived protein adducts in dynamic adduction analyses. **e**, Violin plots of L/H ratios determined from four types of aONE-derived protein adducts in dynamic adduction analyses.

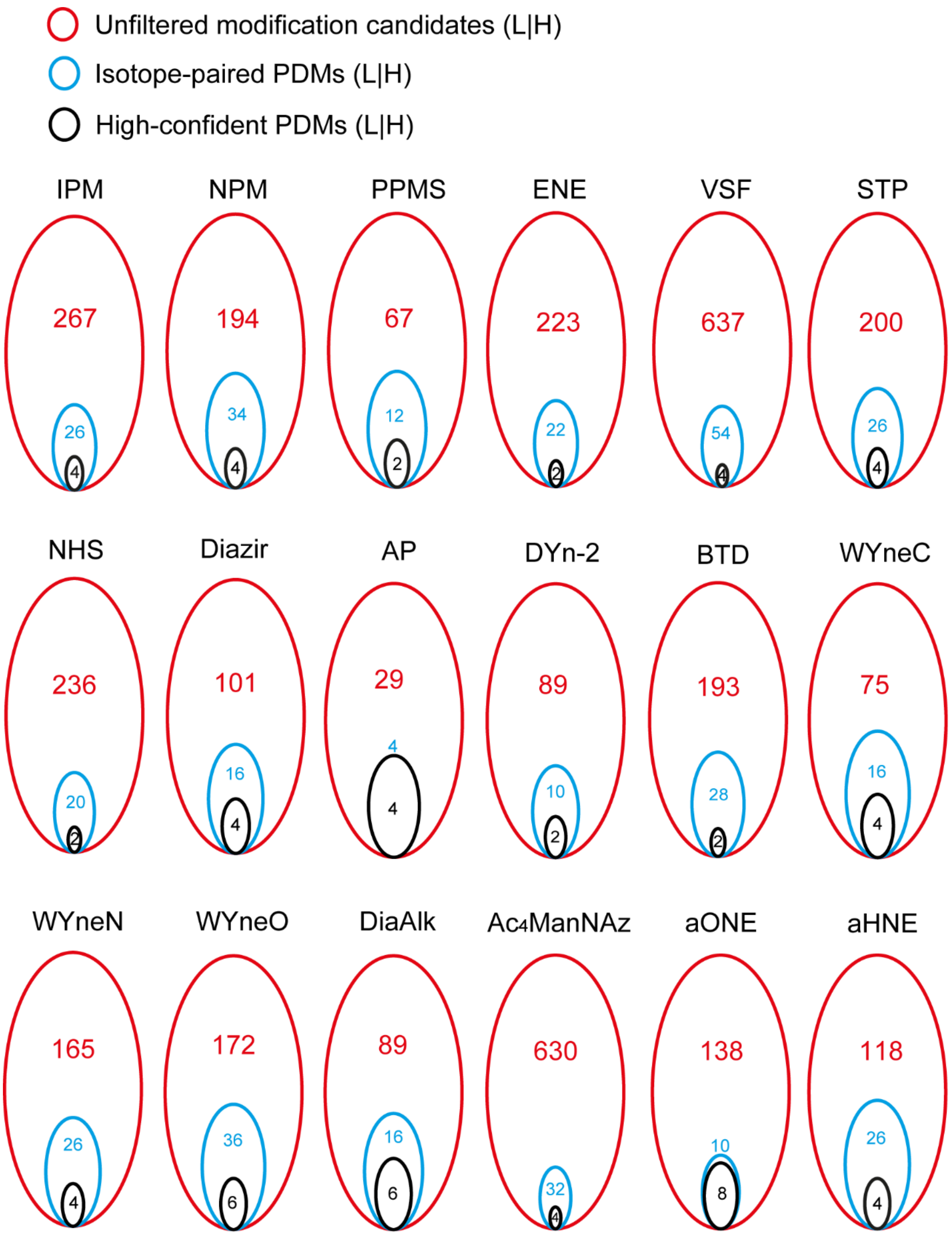


**Supplementary Fig. 15. pChem automatically defines high-confident PDMs.** Overlaps of unfiltered mass shifts higher than 200 Da, isotope-paired PDMs (no cutoffs for L/H ∆mass tolerance and PSM counts) and high-confident PDMs (L/H ∆mass tolerance ≤ 0.001Da, %PSM ≥ 5%).

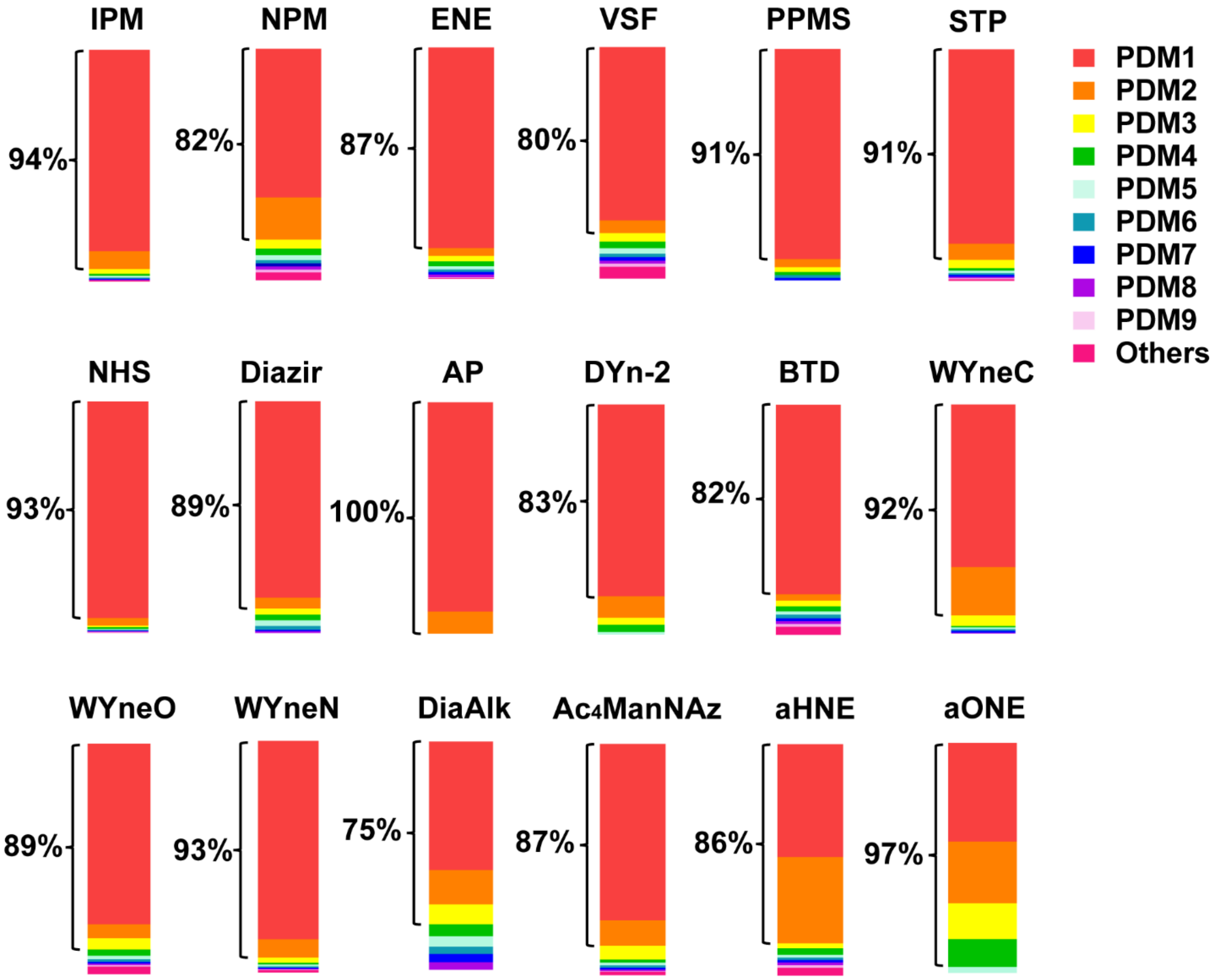


**Supplementary Fig. 16. The predominance of high-confident PDMs.** Stack column plots showing that, for all probes tested herein, the PSMs of the corresponding high-confident PDMs (L/H $\Delta$mass tolerance $\leq$ 0.001Da, %PSM $\geq$ 5%) account for 88.4$\pm$6.3% of those of all isotope-paired PDMs (no cutoffs for L/H $\Delta$mass tolerance and PSM counts).

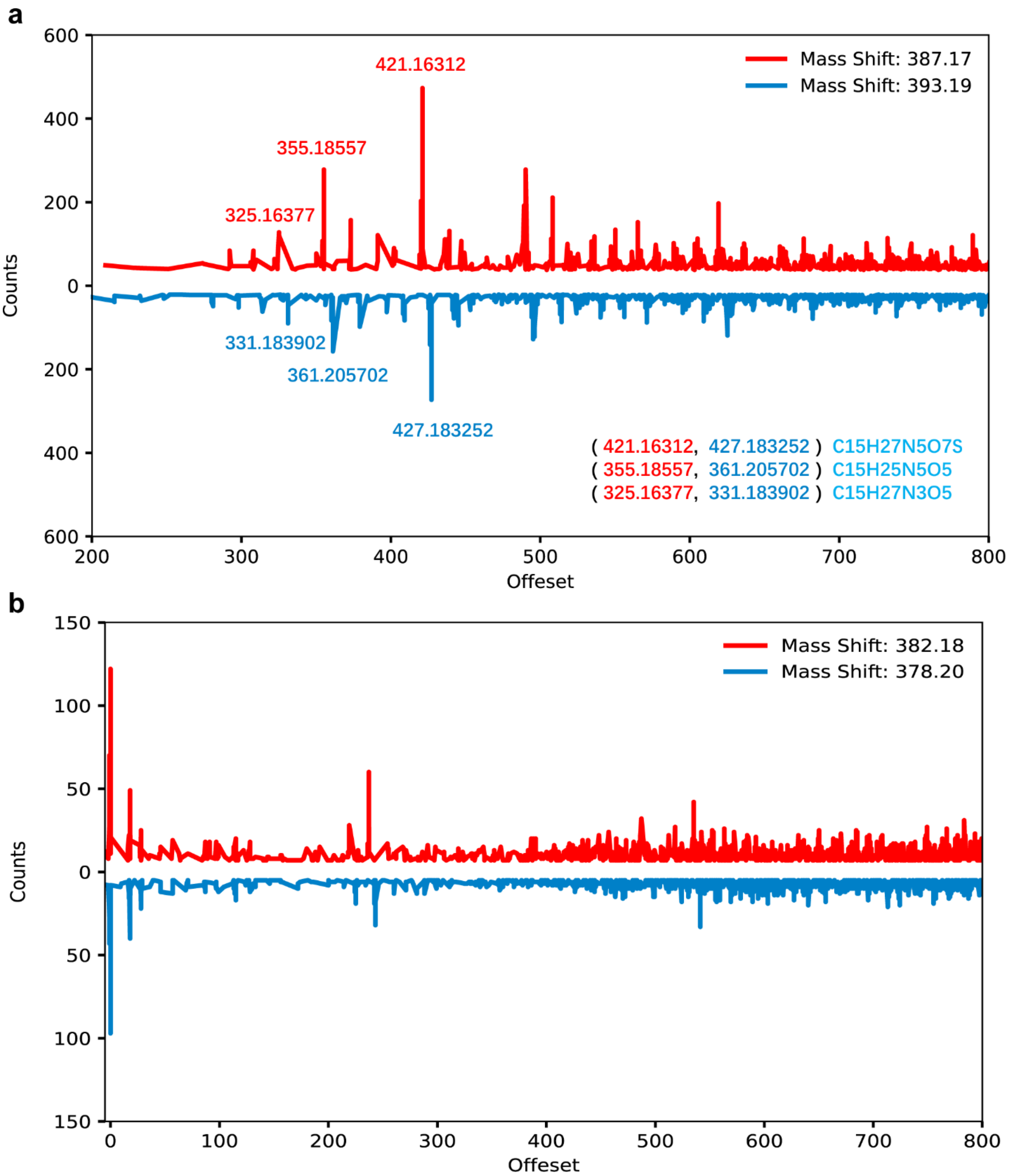


**Supplementary Fig. 17. pChem enables automatic recognition of diagnostic fragment losses (DFLs), if any, for every PDM. a**, Mass offset distribution of the potential DFLs from the DiaAlk dataset. The high-frequency offsets annotated are corresponded to the real neutral losses verified previously. **b**, Mass offset distribution of the potential neutral losses from the AP dataset. The offset with the top-1 frequency is zero, which implicates that the current modification is unlikely to generate any neutral losses.
