## Supplementary Table 1 for "pChem: a modification-centric assessment tool for the performance of chemoproteomic probes"

**Supplementary Table 1. All PDMs identified in this study.**

| Probe | Chemical structures | Probe-derived modifications (PDMs) | Site | Molecular formula of PDM | Theoretical mass of PDM | Notes |
|---|---|---|---|---|---|---|
| **IPM** | 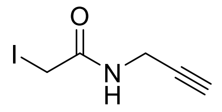 | 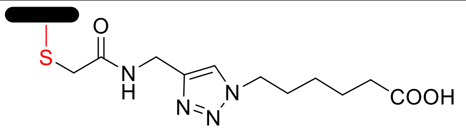 | C | $C_{11}H_{16}N_4O_3$ | 252.12224 | Targeted PDM |
| **NPM** | 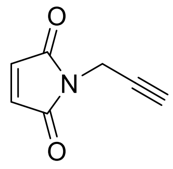 | 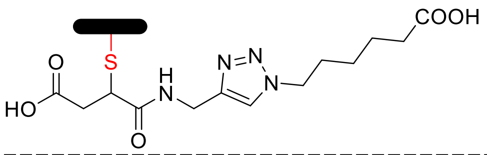 | C | $C_{13}H_{16}N_4O_4$ | 310.12772 | Targeted PDM |
| | | 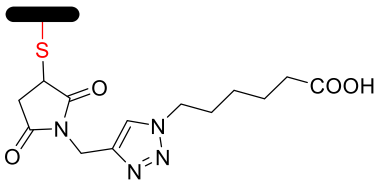 | C | $C_{13}H_{18}N_4O_5$ | 292.11716 | Targeted PDM |
| | | 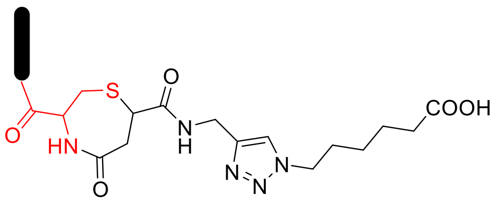 | N-Term | $C_{13}H_{16}N_4O_4$ | 292.11716 | Newly identified PDM |
| **ENE** | 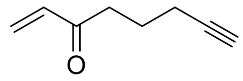 | 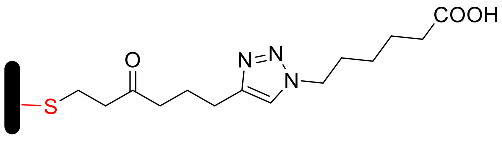 | C | $C_{14}H_{21}N_3O_3$ | 279.15829 | Targeted PDM |
| **VSF** | 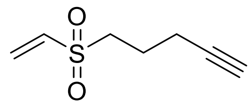 | 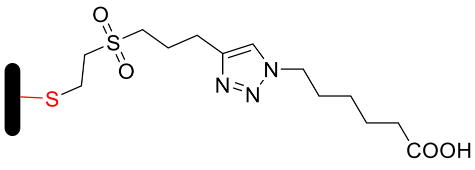 | C | $C_{13}H_{21}N_3O_4S$ | 315.12528 | Targeted PDM |
| **PPMS** | 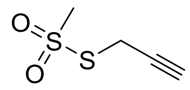 | 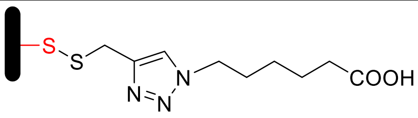 | C | $C_9H_{13}N_3O_2S$ | 227.07285 | Targeted PDM |
| **STP** | 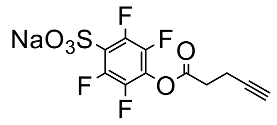 | 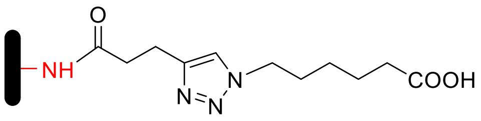 | K | $C_{11}H_{15}N_3O_3$ | 237.11134 | Targeted PDM |
| **NHS** | 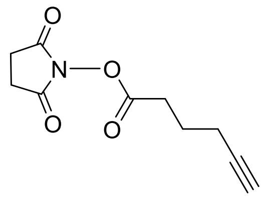 | 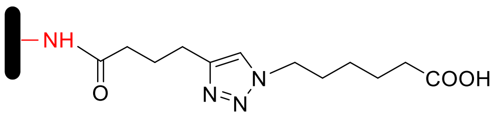 | K | $C_{12}H_{17}N_3O_3$ | 251.12699 | Targeted PDM |
| **Diazirine** | 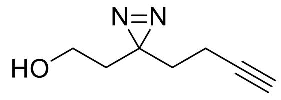 | 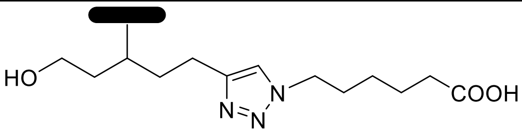 | E | $C_{13}H_{21}N_3O_3$ | 267.15829 | Targeted PDM |
| **AP** | 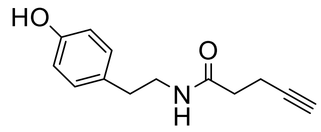 | 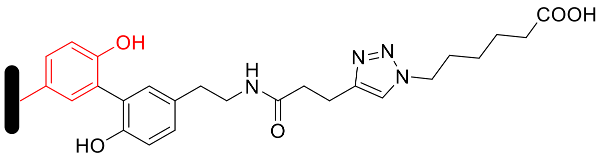 | Y | $C_{19}H_{24}N_4O_4$ | 372.17976 | APEX/HRP |
| | | 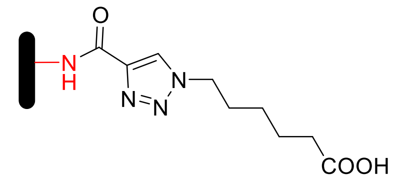 | K | $C_9H_{11}N_3O_3$ | 209.08004 | APEX/Newly identified |
| | | 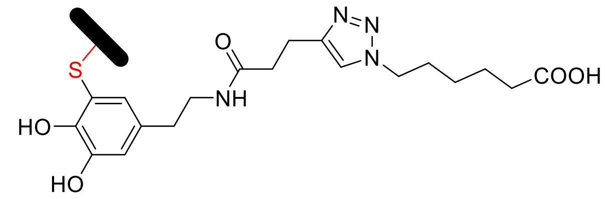 | C | $C_{19}H_{24}N_4O_5$ | 388.17467 | HRP/Newly identified |

| | | | | | | |
|---|---|---|---|---|---|---|
| **DYn-2** | 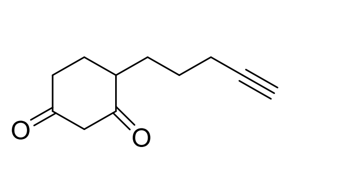 | 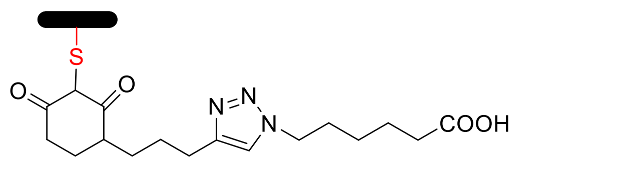 | C | $C_{17}H_{23}N_3O_4$ | 333.16886 | Targeted PDM |
| **BTD** | 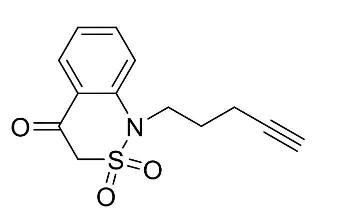 | 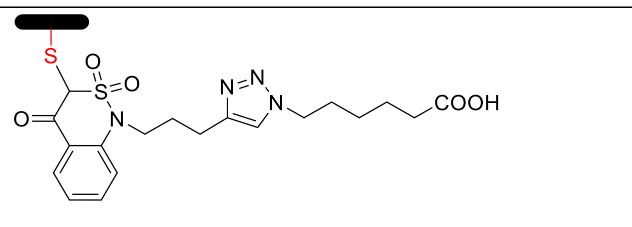 | C | $C_{19}H_{22}N_4O_5S$ | 418.13109 | Targeted PDM |
| **WyneC** | 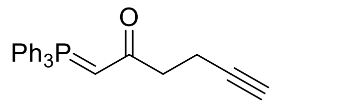 | 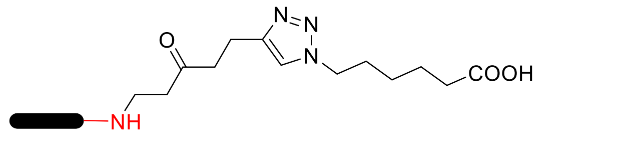 | N-Term | $C_{13}H_{19}N_3O_3$ | 265.14264 | Newly identified PDM |
| | | 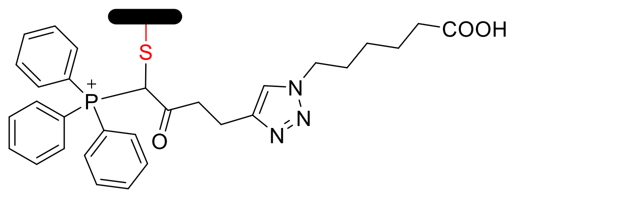 | C | $C_{30}H_{30}N_3O_3P^+$ | 511.20193 | Targeted PDM |
| **WyneN** | 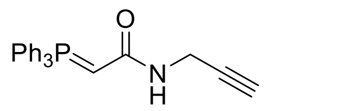 | 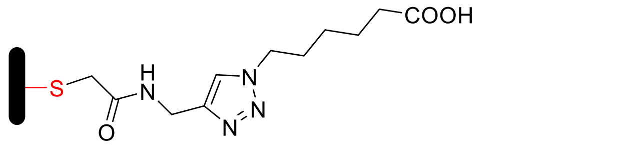 | C | $C_{11}H_{16}N_4O_3$ | 252.12224 | Targeted PDM |
| **WyneO** | 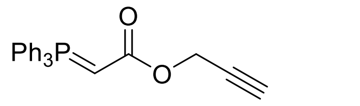 | 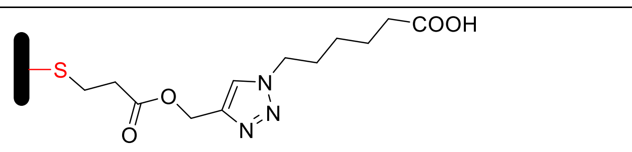 | C | $C_{12}H_{17}N_3O_4$ | 267.12191 | Newly identified PDM |
| | | 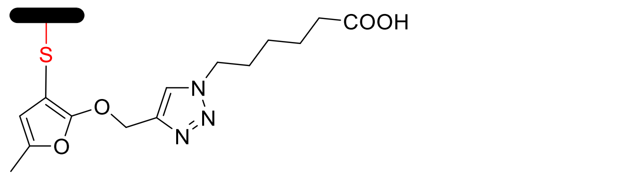 | C | $C_{14}H_{17}N_3O_4$ | 291.12191 | Newly identified PDM |
| **DiaAlk** | 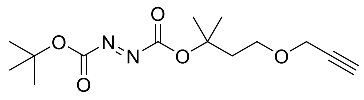 | 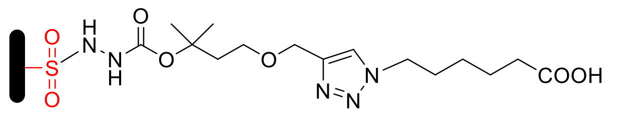 | C | $C_{15}H_{25}N_5O_7$ | 387.17540 | Targeted PDM |
| | | 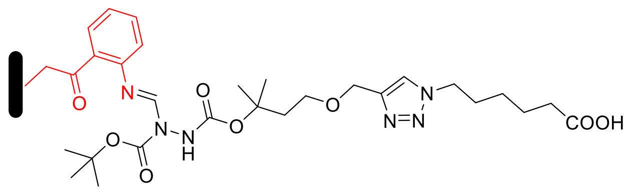 | W | $C_{20}H_{33}N_5O_8$ | 471.23291 | Newly identified PDM |
| | | 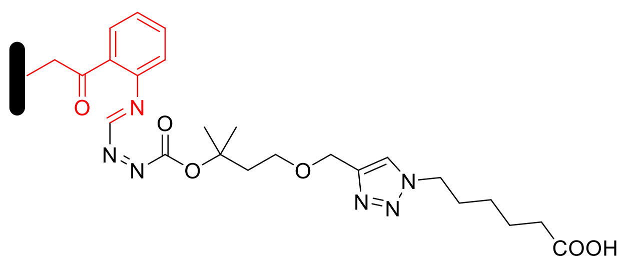 | W | $C_{15}H_{23}N_5O_6$ | 369.16483 | Newly identified PDM |
| **$Ac_4ManNAz$** | 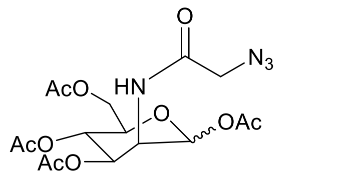 | 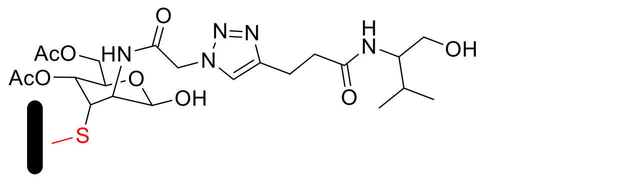 | C | $C_{22}H_{33}N_5O_9$ | 511.22783 | Targeted PDM |
| | | 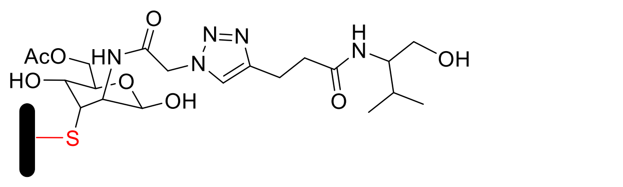 | C | $H_{31}C_{20}N_5O_8$ | 469.21726 | Targeted PDM |

| | | | | | | |
|---|---|---|---|---|---|---|
| **aHNE** | 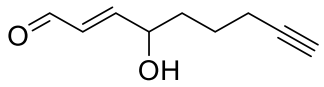 | 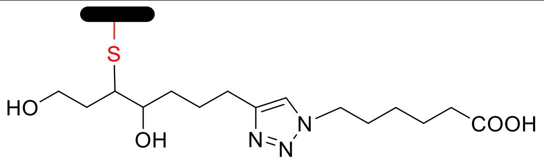 | C | $H_{25}C_{15}N_3O_4$ | 311.18451 | Targeted PDM |
| | | 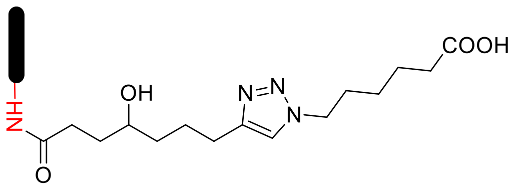 | N-Term | $H_{21}C_{15}N_3O_4$ | 307.15321 | Newly identified PDM |
| **aONE** | 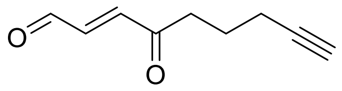 | 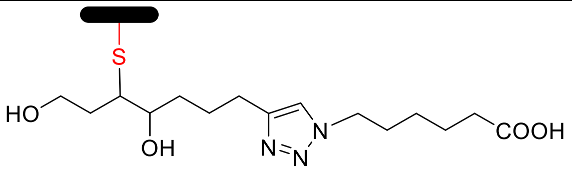 | C | $H_{25}C_{15}N_3O_4$ | 311.18451 | Not detected |
| | | 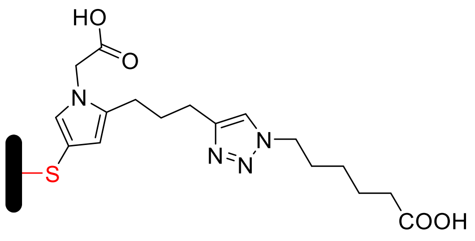 | C | $H_{22}C_{17}N_4O_4$ | 346.16411 | Targeted PDM |
| | | 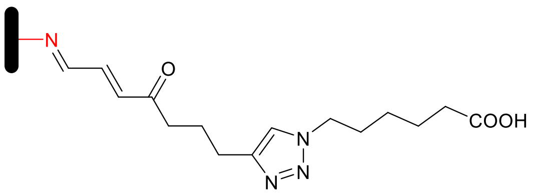 | K | $H_{19}C_{15}N_3O_4$ | 289.14264 | Targeted PDM |
| | | 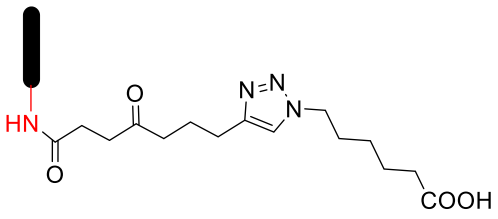 | K | $H_{21}C_{15}N_3O_4$ | 307.15321 | Targeted PDM |
| | | 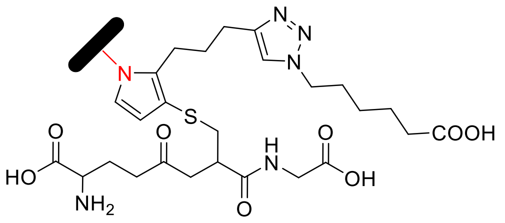 | K | $H_{36}C_{26}N_5O_8S$ | 578.22846 | Newly identified PDM |
